# “Dimerization of the Mineralocorticoid Receptor Ligand Binding Domain by helix 9, 10 and the F-domain”

**DOI:** 10.1101/2020.11.05.369587

**Authors:** Laurent Bianchetti, Deniz Sinar, Camille Depenveiller, Annick Dejaegere

**Author notes:** corresponding author, (AD).

## Abstract

In vertebrates, the mineralocorticoid receptor (MR) is a steroid-activated nuclear receptor (NR) that plays essential roles in water-electrolyte balance and blood pressure homeostasis. It belongs to the group of oxo-steroidian NRs, together with the glucocorticoid (GR), progesterone (PR), and androgen (AR) receptors. Classically, these oxo-steroidian NRs homodimerize and bind to specific genomic sequences to activate gene expression. NRs are multi-domain proteins, and dimerization is mediated by both the DNA (DBD) and ligand binding (LBD) domains, with the latter thought to provide the largest dimerization interface. However, at the structural level, the LBD dimerization of oxo-steroidian receptors has remained largely a matter of debate. This is linked to the receptor refractory expression, purification and crystallization. As a result, there is currently no consensus on a common homodimer assembly across the 4 receptors, *i.e*. GR, PR, AR and MR, despite their sequence homology. Examining the available MR LBD crystals and using widely plebiscited tools such as PISA, PRISM and EPPIC, and the MM/PBSA method, we have determined that an interface mediated by the helices H9 and H10 of the LBD as well as by the F domain presents the features of a biological protein-protein interaction surface. This interface which has been observed in both GRα and MR crystals, distinguished itself among other contacts and provided for the first time a homodimer architecture that is common to both oxo-steroidian receptors.

## 1 Introduction

The mineralocorticoid receptor belongs to the super-family of nuclear receptors (NRs), a metazoan specific super-family of ligand-activated transcription factors that transduce extracellular signals into transcriptional responses. In human, pre- and post-genomic studies have identified 48 NRs that cluster in 7 phylogenetic families (from NR0 to NR6)^1–4^. In the family of steroid hormone receptors (NR3), the oxo-steroidians constitute a group of 4 closely related homologues that bind cholesterol-derived molecules, *i.e*. the progesterone (PR), androgen (AR), mineralocorticoid (MR) and the glucocorticoid alpha (GRα) receptors. From N-to C-terminus, oxo-steroidian receptor sequences exhibit a large N-terminal transactivation domain (NTD), a highly conserved DNA binding domain (DBD), a short hinge region, a ligand binding domain (LBD) and a 10 residue F-domain whose function has remained elusive. The LBD folds into a 12 α-helix sandwich that leads to the formation of a hydrophobic ligand binding pocket (Supp info S1). This modular architecture is a general feature of the NR protein family.

The main regulatory mechanism of NRs is through the binding of a specific ligand in their LBD pocket. For oxo-steroidian receptors, ligand binding is coupled to nuclear translocation and binding to genomic sequences to activate the transcription of target genes. Oxo-steroidian NRs assemble on DNA as homodimers^5–9^. However, higher oligomeric structures such as tetramers have also been reported^10,11^.

Ligand bound NRs adopt a conformation that allows subsequent formation of complexes with transcriptional co-activators, for example co-activators of the steroid receptor coactivator family such as SRC-1, through recognition of a conserved LXXLL sequence motif. NR interaction networks are not limited to co-activators, and much current effort aims at deciphering the complex protein interaction networks associated with NRs, where a partner protein alters gene transcription and enables communication between signaling pathways^12–14^. Reference examples of such regulatory cross-talks are the interaction between GRα and activator protein 1 (AP-1) or nuclear factor κB (NFκB) transcription factors^15,16^.

The interaction surface for the canonical LXXLL co-activator motif is well characterized structurally and uses helices H3, H4, H5 together with the regulatory AF-2 helix H12 to form the docking platform for co-activator proteins^17–19^. Interactions with the peptide motif of co-repressor proteins has also been well described^18,20^. For the oxo-steroidian receptors, interaction surfaces with other proteins, including NR dimer architecture have however remained more elusive. For many NRs, the interaction surface that mediates homodimerization, or heterodimerization with the retinoid X receptor as a partner, has been characterized for both the DBD and LBD^21–23^. In particular, the LBD has been shown in numerous examples to present the largest homo- or heterodimerization interface, implicating helix 9 (H9), 10 (H10) and 11 (H11) (Supp info S1)^23,24^ On the contrary, oxo-steroidian receptors have been reported to use alternative LBD surfaces to homodimerize as for PR^25^, GRα^26,27^ and AR^28^. Functional heterodimeric interactions involving oxo-steroidian receptors have also been documented *e.g*. the peroxisome proliferator activated receptor-alpha (PPARα) and GRα^29^, the AR and GRα^30^ and the GRα and MR^31–36^. However, the structural assemblies for these proteins have remained elusive.

In oxo-steroidian receptors, it would therefore be beneficial to identify interaction surfaces, particularly in the LBD, that could mediate the receptor dimerization and/or interaction with partner proteins. Contacts that are observed in experimental crystal structures can be a rich source of information on functional interfaces. We recently analyzed GRα LBD contacts in crystals^27^. The canonical LBD homodimer implicating helix 9 (H9), 10 (H10) and 11 (H11) was not observed in any GRα LBD crystals, due to the presence of the GRα C-terminal F-domain forming a steric obstacle to its formation^25,27,28,37^. However, other dimeric assemblies of GRα were observed in the experimental X-ray structures^27^, including the architecture proposed as physiologically relevant^26^ (which we refer to as *bat-like*^27^) as well as alternative assemblies not previously discussed. An assembly that was particularly observed was one where the helices 9 were in anti-parallel orientation and the F-domain were at the dimerization interface – referred to in our previous work as the *apH9* complex –^27^. Interestingly, this assembly showed the features of a biological protein-protein interaction^38^, *i.e* significant favorable binding free energy and over-representation of conserved residues; the other observed assemblies did not meet these criteria to the same extent^27^.

In order to further explore the relevance of this interface within the oxo-steroidian group of NRs, we set out to analyze the crystal assemblies of the mineralocorticoid receptor (MR). In the NR superfamily, MR is the closest homologue of GRα (identity 56 % and similarity 75.4 % between human LBD sequences). While GRα is ubiquitously expressed^39^ and controls key physiological processes such as development, response to stress, circadian rhythm, metabolism and homeostasis^40^, MR exhibits a more restricted expression profile. In the kidney and colon, it controls sodium reabsorption and potassium efflux and is involved in electrolyte and blood pressure homeostasis^41^. In human, cortisol and aldosterone are the endogenous ligands of GRα and MR, respectively, and both are secreted by the adrenal cortex, *i.e*. the outer cell layers of the adrenal glands which are located above the kidneys. The structure of the MR DBD homodimer in complex with DNA was resolved by X-ray crystallography^7^ and is similar to that of GRα^42^ as measured by a structural root-mean-square difference of 0.55 Å (PDB-ID:4TNT and PDB-ID:1GLU for MR and GRα DBDs, respectively). In addition, the MR LBD structure was resolved in complex with aldosterone^43^ and dexamethasone^44,45^. Finally, as for GRα, the C-terminus extremity, or F-domain, is a short 10 residue sequence stabilized by a β-strand (S4) and a loop that is packed against the LBD. While the F-domain of oxo-steroidian receptors has been observed to form a steric obstacle to the canonical LBD homodimer formation^25,27,28,37^, it has also been shown to be important for ligand binding and receptor activation in MR^46^ and in GRα^47^.

In the case of MR, no LBD homodimeric assembly has been characterized yet, although several LBD crystals have been deposited (see Table 1)^43,45,48^. In this study, we explored all available MR LBD crystals for protein-protein contacts. In total, 28 crystals were collected from the Protein Data Bank (PDB)^49^ Using protein-protein interaction analysis tools, such as Proteins, Interfaces, Structures and Assemblies (PISA)^50^, Protein-Protein Interaction Prediction by Structural Matching (PRISM)^51,52^ and Evolutionary Protein-Protein Interface Classifier (EPPIC)^53^, several MR-LBD assemblies were determined as being potentially relevant. As expected from the presence of the F-domain, the canonical homodimer was absent in the 28 MR crystals. Moreover, the assembly proposed as physiological for the GRα LBD homodimer^26^, *i.e*. an assembly that brings the loop between H1 and H3 and the C-terminus extremity of H5 at the dimerization interface, was also absent in all MR crystals. In the present study, a MR LBD homodimer mediated by H9, H10 and the F-domain, which shows structural similarity with the GRα *apH9* complex was frequently observed in MR crystals. Using PISA, PRISM, EPPIC, and the Molecular Mechanics Poisson-Boltzmann Surface Area method (MM-PBSA) to calculate binding free energy^54,55^, we explored the biological relevance of this identified MR LBD assembly and compared it to a docked assembly based on the architecture previously identified for GRα *apH9* complex^27^. Although both architectures engage similar interaction surfaces, the docked assembly based on GRα appears more favored at both thermodynamic and residue conservation levels.

**Table 1:**
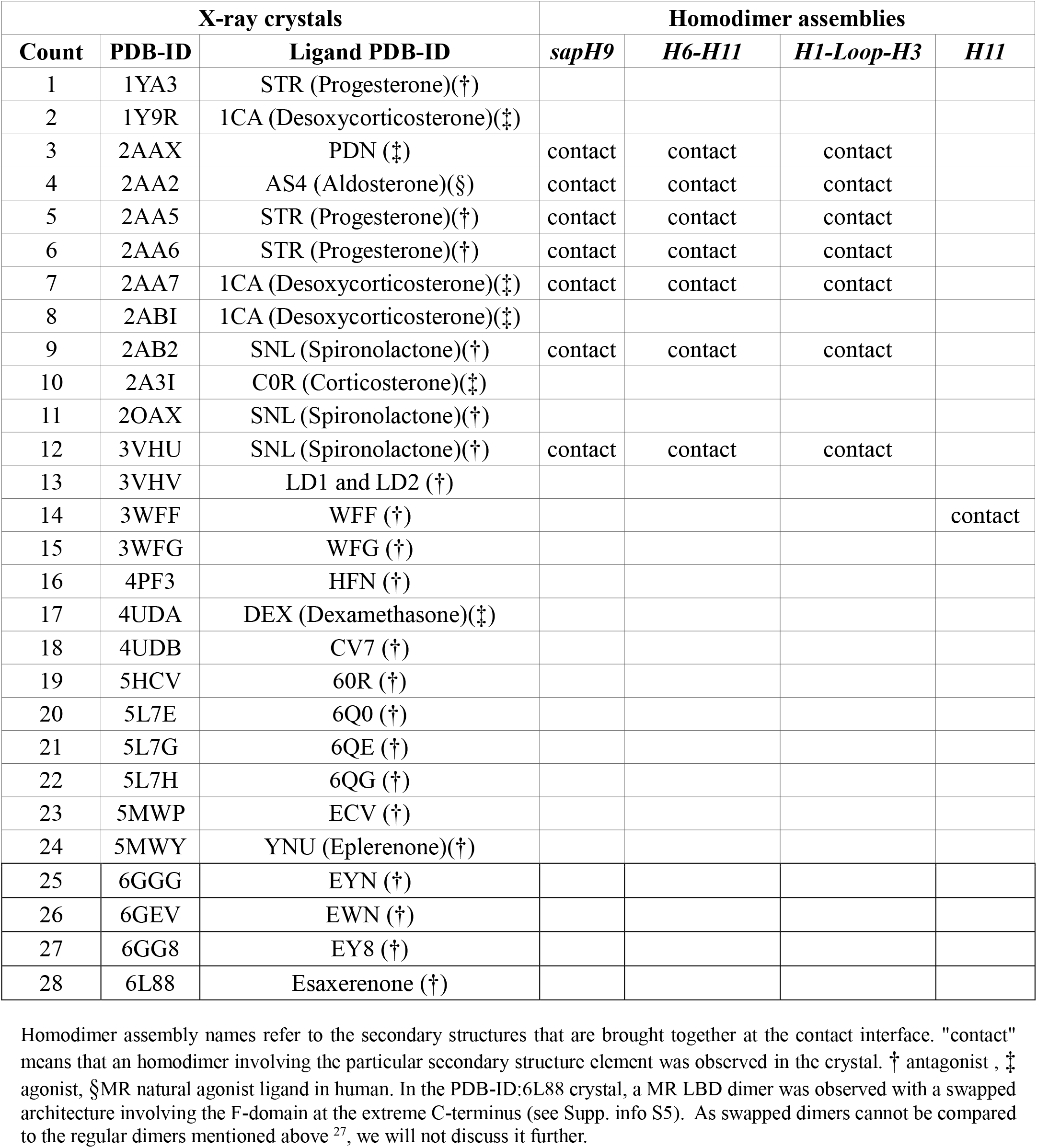
MR LBD structures (X-ray crystals) deposited in the PDB and homodimer assemblies.

## 2 Material and Methods

### 2.1 MR LBD structures

All MR LBD structures were collected from the PDB^49^ using a BLASTP sequence similarity search on the NCBI web server (http://www.ncbi.nlm.nih.gov) with the human MR LBD (RefSeq-ID:NP_000892 from residue Ser 737 to Ser 973) and the F-domain (RefSeq-ID:NP_000892 from residue Gly 974 to Lys 984) sequence as a query with default parameters.

### 2.2 Protein Structural Statistics

MR structures were processed with the protein structural statistics (PSS) tool^56^ to examine the LBD folds in crystals. Receptor chains were structurally superposed using Modeller^57^ and a structurebased multiple sequence alignment was generated. Root mean square fluctuation (RMSF) between the different structures and average B-factor along the LBD sequence were calculated and represented on graphs with gnuplot scripts. To determine the residues that were either structurally unresolved or mutated, the human wild-type MR sequence (RefSeq-ID:NP_000892) was added to PSS sequence alignment and displayed in Aliview^58^.

### 2.3 Bioinformatics tools for protein-protein contact analysis

We used 3 tools to search for protein-protein contacts in MR LBD crystals. First, PDB entries were processed with the PISA program^50^ which calculates interface area (difference in the total accessible surface areas of the isolated monomers and that of the dimeric assembly, divided by 2) and different components of the binding free energy, including the solvation free energy gain upon formation of the interface (ΔiG, kcal/mol). Second, PRISM^51^ was applied to each MR PDB structure to model LBD homodimer assemblies using a precomputed database of template protein interfaces^52^ PRISM constructs assemblies by docking query structural chains on its template complexes and estimates binding free energy. Finally, the EPPIC tool^53^ was used to predict the biological likelihood of homodimeric assemblies using counts of buried residues at the contact interface and scores of amino-acid conservation. First, EPPIC detects all protein-protein contacts in crystals. Second, it counts residues that bury at least 95% of their surface at the contact interface (95%-core-residues). The greater the count of 95%-core-residues, the more likely is the biological relevance of the assembly. Third, enrichment of conserved residues at the contact interface, *i.e*. core-residues that bury a 70% surface threshold (70%-core-residues) *versus* randomly sampled residues on the surface of the complex is calculated using a Z-score. To be significant, the Z-score must be less than −1 which means the 70%-core-residues are enriched in conserved amino-acids.

### 2.4 *sapH9* and *apH9* LBD homodimer models

For both human MR and GRα monomeric LBDs, the available crystal structures present mutations and/or missing residues (Supp info S2). We therefore used modelling tools to obtain a complete 3D structure of the full LBD wild-type sequence. In a first MR model, chain A from PDB-ID:4UDA entry, *i.e*. the LBD bound to dexamethasone, where the structure was determined by X-ray crystallography to 2.03 Å resolution, was selected as a starting structure. Mutated residues were reverted to wild-type amino-acids using the PyMOL Open-Source 1.8.x program^59^. In PDB-ID:4UDA, the loop between H9 and H10 (Lys 909 to Gly 915) and the C-terminal Lys 984 were not resolved. However, residues Lys 909 to Gly 915 were resolved in PDB-ID:3WFF structure (2.05 Å resolution). PDB-ID:4UDA and PDB-ID:3WFF structures were superposed using PyMOL (RMSD = 0.5 Å), and the Lys 909 to Gly 915 atom coordinates were copied from the latter to the former structure while the C-terminal Lys 984 atoms were taken from PDB-ID:2A3I. In a second model, chain A of PDB-ID:2AAX (1.75 Å) was used as a starting structure. The sequence was mutated at position 808 and 810 (Supp info S2) (both residues are located in H5) while atom coordinates of Pro 911 (located in the loop between H9 and H10) and Lys 984 (C-terminus last residue) were unresolved. Both mutated residues were reverted to the wildtype amino-acids and residues Pro 911 and Lys 984 were modelled using the Modeller 9.16 program^57^. The Modeller *automodel and loop-refinement* modules were used and the model with the lowest DOPE score was selected for further analysis. In most MR LBD X-ray crystals, the loop between H9 and H10 is not resolved. Because the H9-H10 loop conformation might play a role in MR complex stabilization, 2 alternative loop conformations observed in crystals were modeled into the structures and used for the binding free energy calculations (Supp info S3 a, b and c). In a first conformation, referred to as the “closed” conformation, the residues of the loop were taken from the structure PDB-ID:2AAX^43^ while residues from the 4UDA-3WFF structure were used to model the loop in a second and “open” conformation. These structures were subjected to an energy minimization and then used to obtain an estimate of the total binding free energy calculated by the MM-PBSA method. Taking into account MM-PBSA results, “closed” loop conformation structures were kept for further analysis while “open” loop conformation structures were not analyzed further (see “Results” for details).

For GRα, chain A of PDB-ID:1M2Z, *i.e*. human LBD bound to dexamethasone^26^, was selected as a starting structure. Mutated Ser 602 was reverted to the wild-type Phe 602. PDB-ID:1M2Z did not present any additional mutation. Moreover, the LBD was fully resolved by X-ray crystallography at 2.5 Å.

Two homodimer architectures were built: *apH9*, built from a unique crystal contact observed in GRα PDB-ID:4P6W and *sapH9* (for shifted *apH9*) observed in seven MR crystals, *i.e*. PDB-ID:2AAX, 2AB2, 2AA2, 2AA5, 2AA6, 2AA7 and 3VHU. MR LBD *apH9* and *sapH9* homodimers were built using the wild-type monomers modeled following the protocol described above (both *apH9* and *sapH9* used the “closed loop” conformation). First, seven MR *sapH9* homodimer models were built by superposing two copies of the wild-type MR LBD onto each of the seven MR *sapH9* complexes observed in crystals. To build the MR *apH9* homodimers, the PDB-ID:4P6W (GRα structure), from which the GRα *apH9* homodimer was originally determined (lattice symmetry is required to observe the contact)^27^, was used as a template. Each MR *sapH9* model was used as a starting structure to build a MR *apH9* model using the PDB-ID:4P6W complex as a template. This resulted in seven additional homodimer assemblies. Finally, two more complexes were generated, *i.e*. a GRα *sapH9* homodimer and the crystallographic GRα *apH9* homodimer^27^. The GRα *sapH9* homodimer was built by superposing two copies of the wildtype GRα LBD onto the MR PDB-ID:2AAX receptor chains A and B, respectively. To built the wildtype GRα *apH9* homodimer, the PDB-ID:4P6W was used as an initial structure. In total, 16 complexes were thus built (Supp info S4).

For both MR and GRα homodimers, cofactors and crystallization additives were removed while crystal water molecules were kept. Protonation states of titratable residues were determined at pH 7.4 with the PROPKA program^60^ implemented in the PDB2PQR server^61^. Dexamethasone was built using Avogadro 1.0.3^62^ and the molecular geometry was optimized. Dexamethasone molecules were then placed into the binding pocket of the LBDs by atom alignment with PyMOL using the ligands already resolved in the crystal structures. Force field parameters for dexamethasone were obtained from the PARAMCHEM webserver^63^ and used without further modification. Hydrogen atom placement on the proteins was performed using the HBUILD^64^ facility in the CHARMM program^65^. The potential energies of ligand bound homodimers were minimized with 500 steps of steepest descent (SD) algorithm using CHARMM program version c37b1^65^ with non-bonded interactions truncated at 14 Å distance using switch and shift functions for van der Waals (vdW) and electrostatic forces, respectively.

### 2.5 Molecular dynamics (MD) simulations and binding free energy calculations

As a preliminary analysis, binding free energies of MR and GRα *sapH9* and *apH9* LBD homodimers were calculated on energy minimized assemblies as described above.

However, to take into account structural fluctuations that can impact binding free energy calculations, assemblies were submitted to a 10 ns MD simulation. Molecular trajectories were performed with the NAMD program^66^ using the CHARMM27 all-atom force field^67^. The homodimers were immersed in a cubic box of TIP3P water molecules (110 × 110 × 110 Å volume). Chloride and sodium counter-ions were added to attain neutralization at a physiological concentration of 0.15 M. MD simulations were started with 2 phases of water minimization and heating while atom coordinates of the protein complex were fixed. In the first phase, water was minimized by 1000 steps of conjugate gradient (CG) and heated at 600K. This was followed by a second phase where water was minimized by 250 steps of CG and heated to 300K. Complexes and water molecules were then minimized by 2000 steps of CG and subsequently heated at 300K. Periodic boundary conditions were used and the particle mesh Ewald (PME) algorithm was applied to take into account long-range electrostatic interactions. All bonds between heavy atoms and hydrogens were constrained using the SHAKE algorithm and an integration time step of 1 fs was used for all simulations. The system was equilibrated for 150 ps. This was followed by the production phase yielding a 10 ns MD simulation. In total, 16 MD simulations were thus carried out and the total binding free energies were calculated for each simulation by the MM-PBSA method^54,55^.

An automated procedure based on the MM-PBSA method^55^ was used to obtain the total and per-residue energetic contribution to homodimer formation. The Gibbs binding free energy upon proteins association can be expressed by:

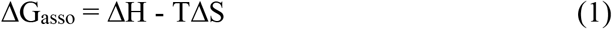

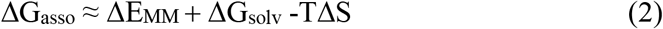

where ΔE_MM_ is the total molecular mechanical energy variation, ΔG_solv_ is the solvation free energy variation and −TΔS is the conformational entropy variation upon complex formation. ΔE_MM_ can be calculated as

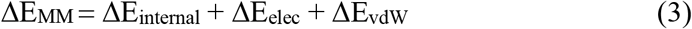

where ΔE_internal_ is the energy variation associated with bond lengths, angles and dihedrals, ΔE_elec_ and ΔE_vdW_ represent electrostatic and vdW terms, respectively.

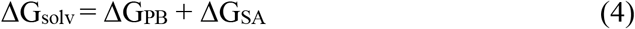

ΔG_PB_ and ΔG_SA_ are solvation energy variation associated with polar and non-polar contributions, respectively. The conformational entropy change was not estimated in this analysis. In addition, ΔE_internal_ was equal to 0 as the monomeric structures were generated simply from the MD simulations of the dimers. There are therefore no changes in internal conformation between the chain in the dimer and the monomeric chain. To identify structures for the MM-PBSA analysis, we computed the Coulomb interaction energy *in vacuo* between the protein chains for all conformations saved from the MD trajectory. A dielectric constant of 1 and a nonbonded cutoff of 12.5 Å were used with a shift truncation function for electrostatics. The conformations of each trajectory were clustered in 10 groups based on their electrostatic interaction energy. The conformation with vacuum interaction energy closest to the cluster average value was extracted and processed using the MM-PBSA procedure. In all, 10 conformations were extracted from molecular dynamics trajectories in this nonlinear fashion. The results for each representative structure were weighed by their respective cluster population and averaged to obtain the total and per residue contribution to the binding free energy.

Finally, Equation (2) can then be written as

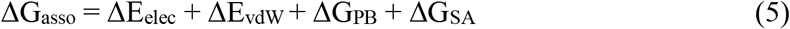

The protein and solvent contribution to the electrostatics term were calculated using the University of Houston Brownian Dynamics (UHBD)^68^ or the Adaptative Poisson-Boltzmann Solver (APBS)^69^ with a grid spacing of 0.3 Å while the vdW and solvent accessible terms were calculated using CHARMM^65^.

## 3 Results

### 3.1 Structural statistics of PDB deposited MR LBD structures

A total of 28 MR LBD crystal structures were exhaustively collected from the PDB (Table 1). All structures were human sequences and bound to a ligand (holo form). The 28 PDB entries contained a total of 47 polypeptide receptor chains that were processed with the protein structural statistics (PSS) program^56^ with the exception of PDB-ID:6L88. Indeed, all structures except PDB-ID:6L88 exhibited a regular LBD fold (Supp info S5 and S6a). Using the superposition of the receptor chains, the C-terminus extremity, *i.e*. the F-domain, showed low root mean square fluctuations (RMSF) between the structures (Supp info S6b) and low average B-factor (Supp info S6c), except for the last few residues. This indicates that it is tightly packed against its self-LBD with low conformational freedom in the X-ray structures. In its observed crystallographic position, the F-domain prevents the formation of the canonical dimerization surface through steric hindrance, in agreement with previous reports^25,27,28,37^. The structural statistics also indicate, as could be expected, low structural fluctuations in elements of secondary structure and larger fluctuations in loops, with the loop between H9 and H10 showing the largest structural variations (Supp info S6b and c).

### 3.2 Assemblies of MR LBD homodimers observed in crystals

Using the PISA program^50^, four homodimeric assemblies were retrieved from the crystal structures and named according to the secondary structures that build up the contact interface (see Table 1). Interestingly, the assemblies observed for MR were for the most part different than those observed for GRα^27^ (Supp. info S7). A first MR assembly brought H9, H10 and the F-domain to the dimerization interface and buried an average of 610 +/- 100 Å^2^ of solvent accessible surface per monomer (Figure 1a). In this architecture, the H9 helices exhibited an anti-parallel orientation and presented structural similarity with the *apH9* - anti-parallel H9 – homodimer reported for GRα (Figure 2a and b)^27^ However, some differences were noticed, *i.e*. the distance between the H9 helices at the dimerization interface was greater in MR than in GRα homodimer while it was the opposite for H10 helices. In addition, H9 of both monomers were face to face in GRα *apH9* but shifted by one helix turn in the MR complex (Figure 2b). Therefore, we named the MR assembly *sapH9* for “shifted *apH9*”. In the crystal structures, all MR *sapH9* contacts showed this H9 helix turn shift (Supp. info S8). This difference in contacts between the MR and GRα H9 and H10 helices is not easily related to the primary sequences of both GRα and MR which are highly similar in this region except for the loop between H9 and H10 (Figure 2c).

**Figure 1:**
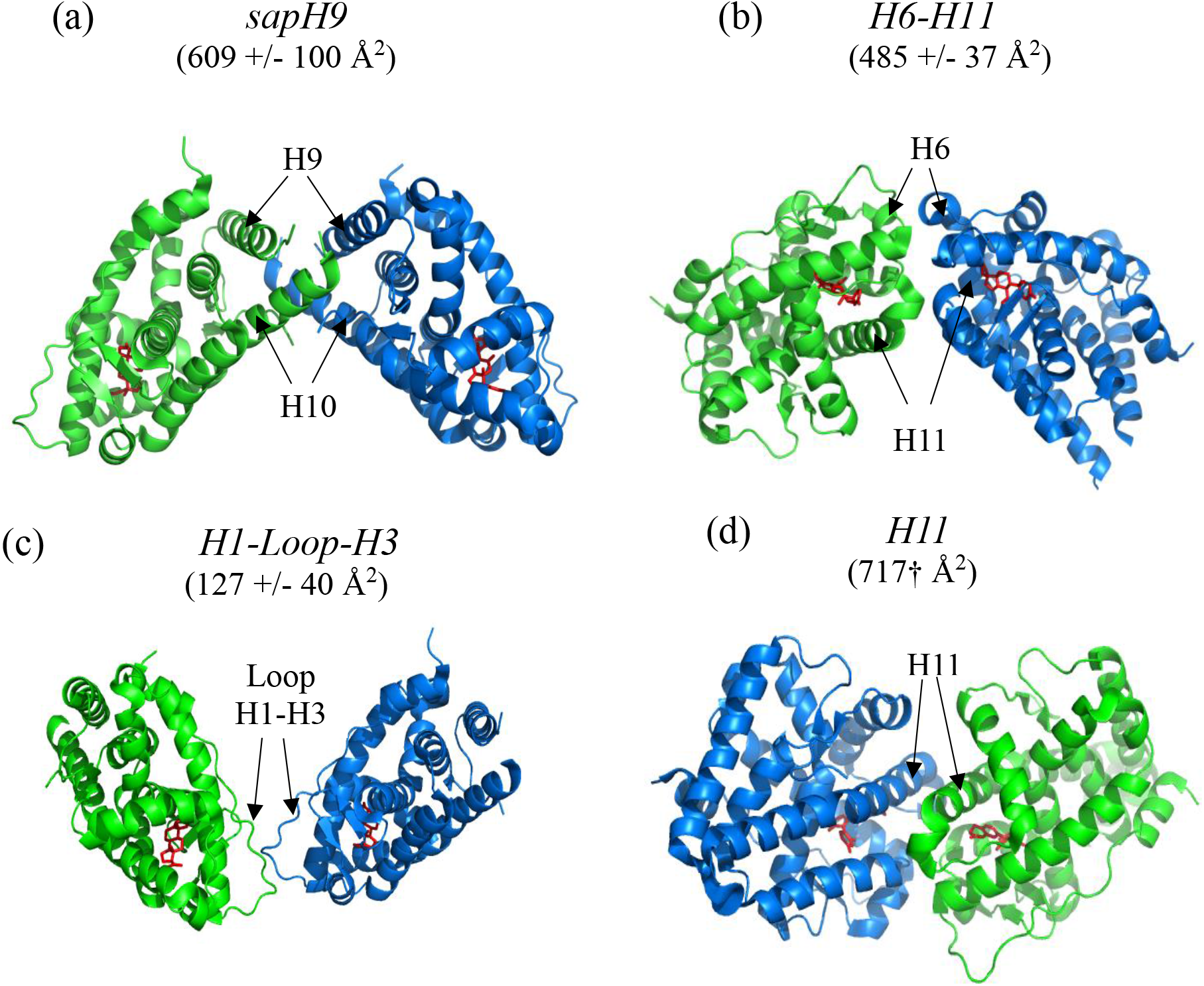
MR LBD homodimeric assemblies observed in crystals. Average buried solvent accessible surface areas per monomer upon forming the dimer are indicated in parenthesis. † symbol means that only one PDB with the shown dimeric contact was available. Ligands are colored in red. Monomers are colored in green and blue.

A second assembly, involving H6 and H11 was called *H6-H11* and buried 485 +/- 37 Å^2^ per monomer (Figure 1b). A third assembly that buried only 127 +/- 40 Å^2^ at the dimerization interface involved the loop between H1 and H3 (*H1-Loop-H3*) (Figure 1c). The *sapH9, H6-H11* and *H1-Loop-H3* assemblies were observed in the same seven PDB entries (Table 1) and they were observed in the presence of both agonist and antagonist ligands. Finally, a fourth homodimeric architecture that brought H11 to the dimer interface was observed only once (named *H11*) and showed a large buried surface *of* 717 Å^2^ per monomer (Figure 1d). Of note, *H6-H11, H1-Loop-H3* and *H11* assemblies were not previously observed in any GRα LBD crystals^27^. Therefore, only assemblies that brought H9 in anti-parallel orientations were simultaneously present in the crystals of both receptors. Since our previous report on GRα homodimerization^27^, eight additional GRα LBD structures have been deposited in the PDB, *i.e*. PDB-ID:5NFP, 5NFT, 5UC1, 5UC3, 6DXK, 6EL6, 6EL7 and 6EL9^70–73^. In 5 crystals out of 7, a new complex architecture was observed that brought the C-terminus of H12 to the contact interface. However, only 157.8+/-14.9 Å^2^ of protein surface was buried per monomer, which is not indicative of a stable interface. In addition, a previously unreported assembly that made contact between monomer by the H7 was observed in PDB-ID:5UC3 and 6DXK. In these latter structures, the H12 showed an atypical position that is not compatible with coactivator binding or was structurally unresolved.

For MR, no contact between regularly folded LBDs was present in the remaining PDB entries. The canonical NR homodimer through helix 9 (H9), 10 (H10) and 11 (H11) was not observed in any of the MR PDB entries. In addition, neither the *bat-like* complex which had been previously reported for GRα LBD^26,27^ nor the frequent H1 assembly^27^ were observed in MR crystals. Finally, only one swapped dimer (Supp. info S5), *i.e*. an assembly that exchanges secondary structures between monomers, was observed for MR while several were observed for GRα^27^.

### 3.3 MR *sapH9* homodimer stability is supported by both PISA and PRISM

To compare the stability of the different dimer architectures, we used estimates of stability provided by PISA and PRISM. Although these estimates should be taken with caution owing to the approximations involved, they provide a ranking of the different complexes. PISA analyses proteinprotein contacts in crystals and provides an estimate of interface stability, related to the solvation free energy gain upon interface formation. PRISM docks query protein structures to a precomputed database of template oligomeric complexes and interfaces and computes a binding energy score for each dimer model. PRISM therefore proposes some architectures that are similar to those observed in crystal contacts of MR, and also new, docked structures, mostly inspired by the contacts observed for GRα. However, PRISM does not systematically propose all the assemblies found for MR in crystal packing, as it depends on its database of template interfaces, which is not exhaustive. We observed that both methods consistently proposed the *sapH9* as the most stable assembly among seven different evaluated complex architectures, *i.e. sapH9, H6-H11, H1-Loop-H3, H11, H1, Bat-like* and *pH11-H12* (see Figure 3a and b and Supp. info S9).

### 3.4 Biological likelihood of MR *sapH9* is supported by EPPIC

MR LBD crystal structures were submitted to the EPPIC program^53^. The criteria used by EPPIC to estimate the biological relevance of a protein-protein assembly are the number buried amino acids at the contact interface and the enrichment in conserved residues at the interface as measured by a Z-score. The method is sensitive to details of the architecture, as related architectures, such as *sapH9* in different PDB entries, obtain different scores. However, out of all architectures examined, the *sapH9* consistently came out as having the largest number of fully buried amino acids, the best interface conservation score, and the best biological likelihood (Figure 3c).

### 3.5 H9-H10 loop is involved in MR *sapH9* stabilization

Based on the results of PISA, PRISM and EPPIC, the *sapH9* assembly was further investigated while the other assemblies were discarded. As noted above, the *sapH9* architecture can have different PISA/PRISM stability estimates and EPPIC assessment that vary as a function of the PDB structure used, even if the architecture is conserved (see Table 1, Figure 3 and Supp. info S9). We investigated whether the position of the loop between H9 and H10 which is implicated in the interface (see Figure 2b) and shows large positional fluctuations (see section 3.1) could be linked to significant energy variations in complex stabilization. Since the residues of the loop were not resolved in a majority of MR LBD structures, the loop is flexible. An MR *sapH9* homodimer based on the conformation of the H9-H10 loop in the “closed” conformation, as seen in PDB-ID:2AAX, exhibited both lower PISA solvation free energy gain (ΔiG = −5.1 kcal/mol) and greater contact surface area (967 Å^2^) than an MR *sapH9* assembly with the H9-H10 loop in the “opened” conformation as seen in PDB-ID:3WFF, *i.e*. PISA solvation free energy gain of −4.2 kcal/mol and 541 Å^2^. This result was confirmed by subjecting this energy minimized MR *sapH9* complexes in both “opened” and “closed” H9-H10 loop conformations to the more rigorous MM-PBSA analysis (Supp. info S3d and e). The binding free energies of both *sapH9* and *apH9* complexes were greater (absolute value) for “closed” loop conformation structures than for “opened” loop conformations. In view of these data, we decided to keep the “closed” loop conformation for further analysis using MM-PBSA methods. This allowed the comparison of the most stable *sapH9* architecture with the previously determined GRα *apH9* based architecture (see below).

**Figure 2:**
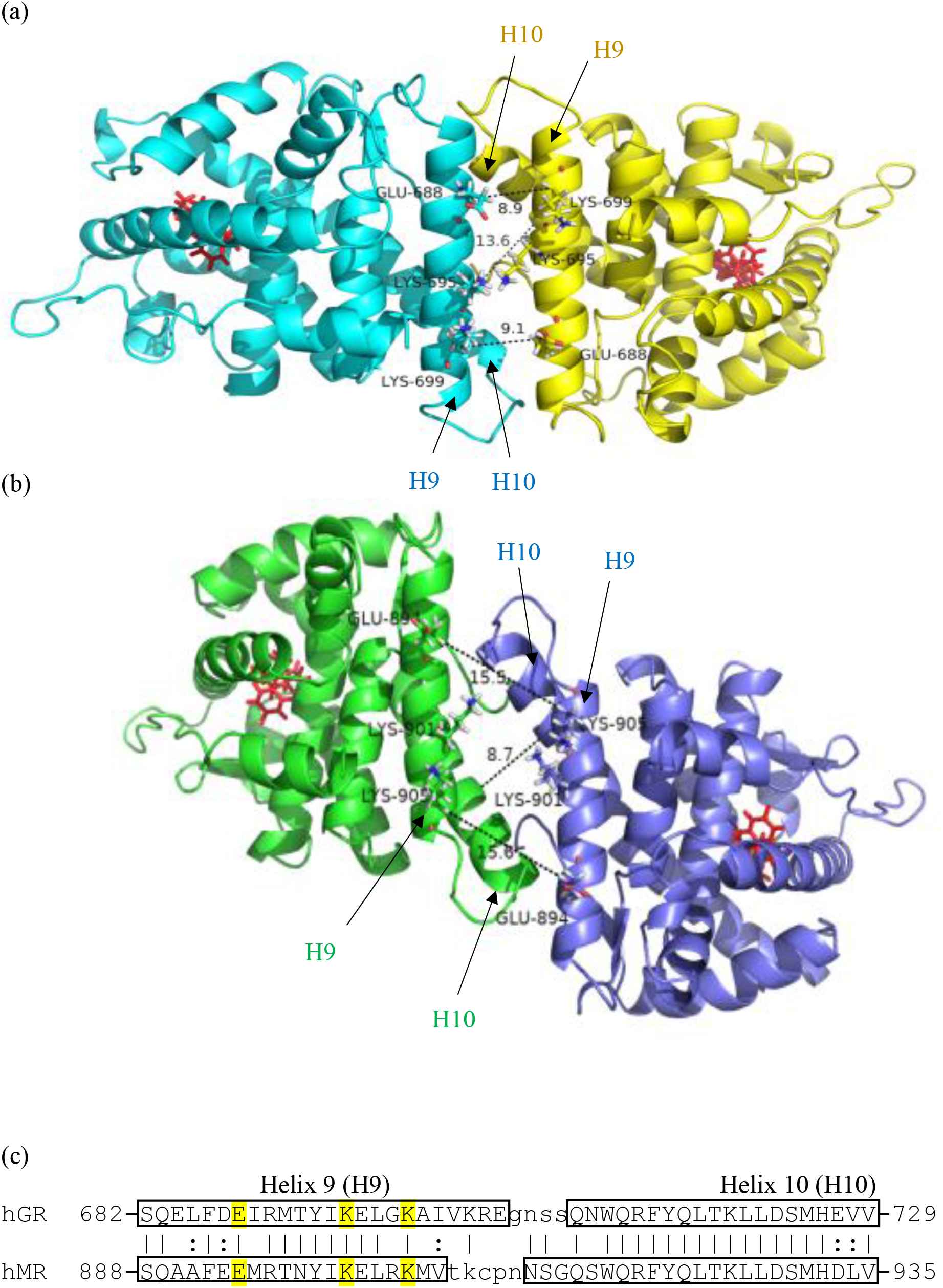
Assemblies that show H9 in anti-parallel orientations at the dimer interface. Ligands are colored in red. Distances are measured in Å and indicated with dashed lines. Distance between the H10 of both monomers was measured perpendicularly to the helices and distance between residues was measured between α-carbons. (a) GRα *apH9* assembly as observed in the crystal of PDB-ID:4P6W. Monomers are colored in yellow and cyan. (b) MR *sapH9* assembly as observed in the crystal structure of PDB-ID:2AAX (Except Pro 911 in the loop between H9 and H10 which was unresolved and therefore structurally modelled). Monomers are colored in green and blue. (c) Human MR (hMR) and human GRα (hGRα) pairwise sequence alignment showing the H9 and H10 region. Residues with yellow background are indicated on figures a and b. Residue identities and similarities are indicated with pipe and colon characters, respectively.

### 3.6 MM-PBSA indicates that the *apH9* architecture is more stable than the *sapH9* assembly

As mentioned above, the crystal protein-protein contacts observed in the *sapH9* MR dimer (see Figure 1) are predicted to be the most relevant identified for MR using a variety of assessment methods (see above). Furthermore, this assembly bears resemblance to an assembly we previously observed as stable for the homologous receptor GRα^27^. In order to see which of the two assemblies, the *sapH9* or the *apH9* is likely to be most stable, we compared four assemblies, *i.e*. MR *sapH9*, MR *apH9*, GRα *sapH9* and GRα *apH9*. Of those, MR *sapH9* and GRα *apH9* were observed as crystal contacts while MR *apH9* and GRα *sapH9* were modelled (see Material and Methods for details).

As a first approach to estimate the stability of complexes, we subjected the structures to an energy minimization followed by the MM-PBSA analysis (Figure 4a and Supp. info S10). We then applied a second approach, which takes into account atom motions. In this second approach, we applied the MM-PBSA analysis on representative structures determined from an MD simulation and averaged the results as described (see Material and Methods for details) (Fig 4b and Supp. info S11). For MR, we used the seven *sapH9* complex structures identified through crystal contacts as starting structures for an MM-PBSA analysis (see Table 1). Moreover, we constructed seven MR *apH9* complexes through modelling. For GRα, we used the sole available complex in *apH9* conformation obtained from PDB-ID:4P6W^27^ The same procedure was applied to the single docked GRα *sapH9* structure. Thus in total, 16 independent complexes were processed.

The results from this analysis (Fig 4a, b and Supp. info S10 and S11) indicate that for the starting structures of both MR and GRα, the *apH9* architecture is more stable than that of *sapH9*. Concerning the binding free energies, the value for the human GRα *apH9* dimer obtained from the simulation was consistent with the value obtained for mouse GRα^27^. In further analysis of the MR assemblies, we calculated the free energy decomposition by the MM-PBSA method using the structures from the MD simulations, as described in Methods. The decomposition determines the amino-acids that make dominant contributions to binding free energy (Supp. info S12), *i.e*. the so-called hotspot residues. For MR, structures PDB-ID:2AAX and PDB-ID:3VHU were selected to represent *sapH9* and *apH9*, respectively, as they showed the most negative total binding free energies for their respective assemblies. For GRα, only one simulation was available. The decomposition showed that certain amino acids of loop 9-10, *i.e*. Ser 708 and Ser 709 in GRα and Thr 908 to Asn 913 in MR, contributed to the stabilization of both the *sapH9* and *apH9* dimers (Supp. info S12), which is coherent with the difference in stability observed as a function of loop conformation for MR *sapH9* (see above 3.5). Of particular note, the H9-H10 loop is moderately flexible and has been fully resolved in all available GRα LBD structures^27^.

However, dominant contributions to the binding free energy are provided by residues from helices H9 and H10. The decomposition analysis showed that Trp 712 in GRα H10 (*i.e*. Trp 918 in MR H10) is buried at the interface in both the *apH9* and *sapH9* architectures and makes a significant contribution to the dimer structural stabilization (Figure 5). This is consistent with data obtained on mouse GRα^27^. Finally, while both *apH9* and *sapH9* assemblies pack hydrophobic residues such as Trp 712, Phe 774 and Met 691 in GRα (Figure 5a, b, e and f) and Trp 918, Leu 979 and Ile 881 in MR (Figure 5c, g and h) at the contact interface, the *apH9* complexes also establish a network of salt-bridges across the dimerization interface, *e.g*. Glu 688 and Lys 695 in GRα (Figure 5b), and Glu 894 and Lys 905 in MR (Figure 5d). This further supports the conclusion that *apH9* is likely to be more stable than *sapH9*.

### 3.7. Biological likelihood of MR *apH9* is greater than MR *sapH9*

Using energy minimized structures, the seven MR LBD *sapH9* and the seven MR LBD *apH9* complexes were submitted to the EPPIC program and bioprobabilities were determined^53^. In addition, both GRα *apH9* and *sapH9* were processed as reference points. The bioprobability of the GRα *apH9* was 0.47 while the GRα *sapH9* was 0.25. Five minimized MR *sapH9* assemblies showed a bioprobability greater than 0.65, while 2 exhibited a bioprobability less than 0.3. On the contrary, all MR *apH9* complexes showed a bioprobability greater than 0.85. A one-sided Wilcoxon test was carried out to determine if bioprobabilities were greater for *apH9* than for *sapH9* MR complexes (after square root arcsinus mathematical transformation) and produced a p-value of 0.0042. Therefore, according to EPPIC, the bioprobability is significantly greater for MR *apH9* than for MR *sapH9* assembly (Figure 3d).

**Figure 3:**
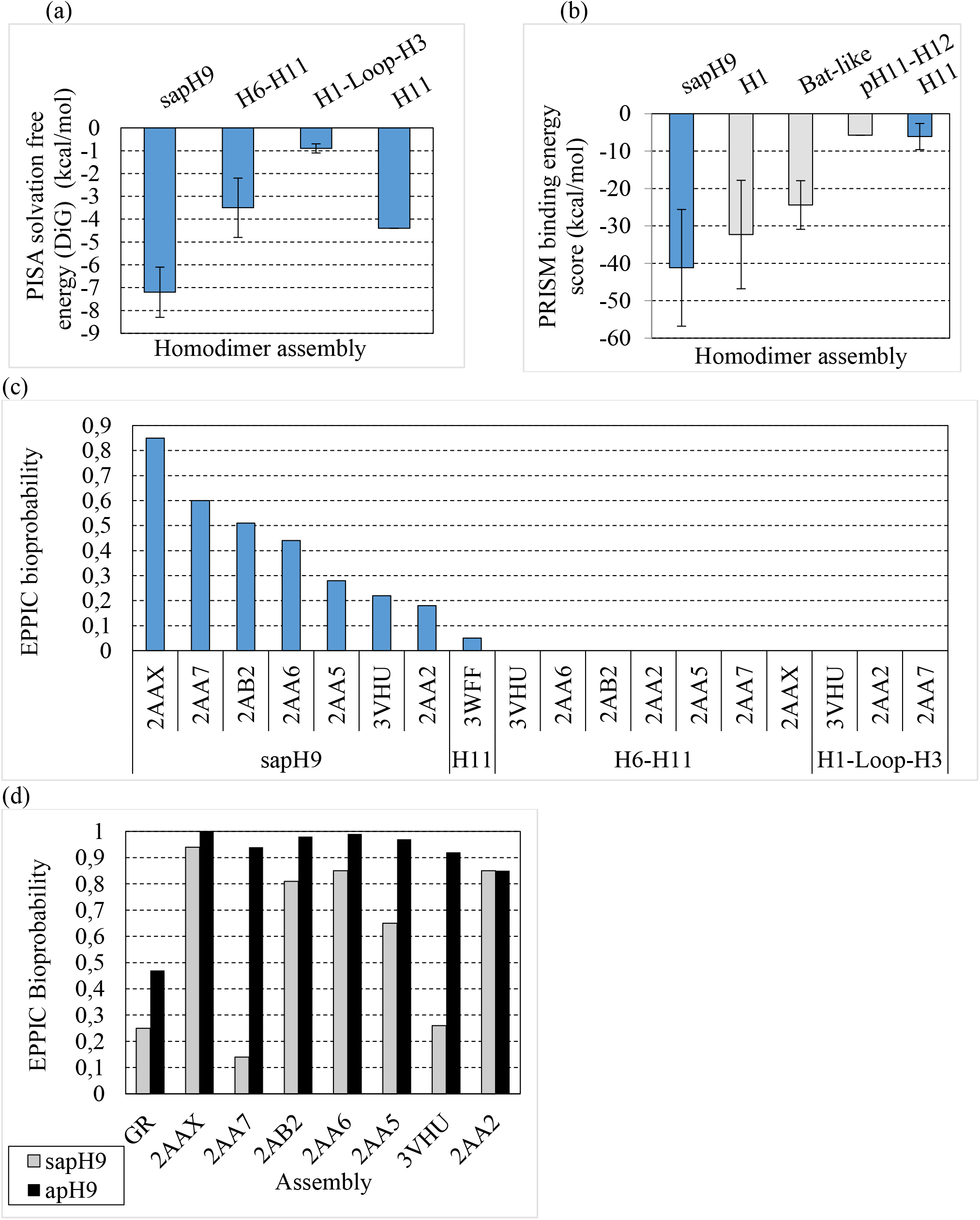
Binding free energy of MR LBD homodimers calculated by bioinformatics tools. (a) PISA solvation free energy gain (Δ^i^G) for the four assemblies listed in Table 1. (b) PRISM binding energy scores. Blue bars correspond to complexes that are based on a template assembly found in MR LBD crystals while gray bars corresponds to complexes where the template is present in a GRα assembly. Of note, not all architectures observed for GRα^27^ or MR correspond to PRISM templates, and therefore the proposed docked complexes do not exhaustively cover the crystal contacts of MR and GRα, but only a subset. In particular, the *apH9* complex of GRα is not present in PRISM database as a template. (c) EPPIC analyses of the MR LBD homodimers (see Table 1), bioprobabilities are calculated using crystal structures (0.5 threshold). (d) Using energy minimized structures, the seven MR LBD *sapH9* and the seven MR LBD *apH9* complexes were submitted to the EPPIC program and bioprobabilities were determined while both GRα *apH9* and *sapH9* were processed as reference points.

**Figure 4:**
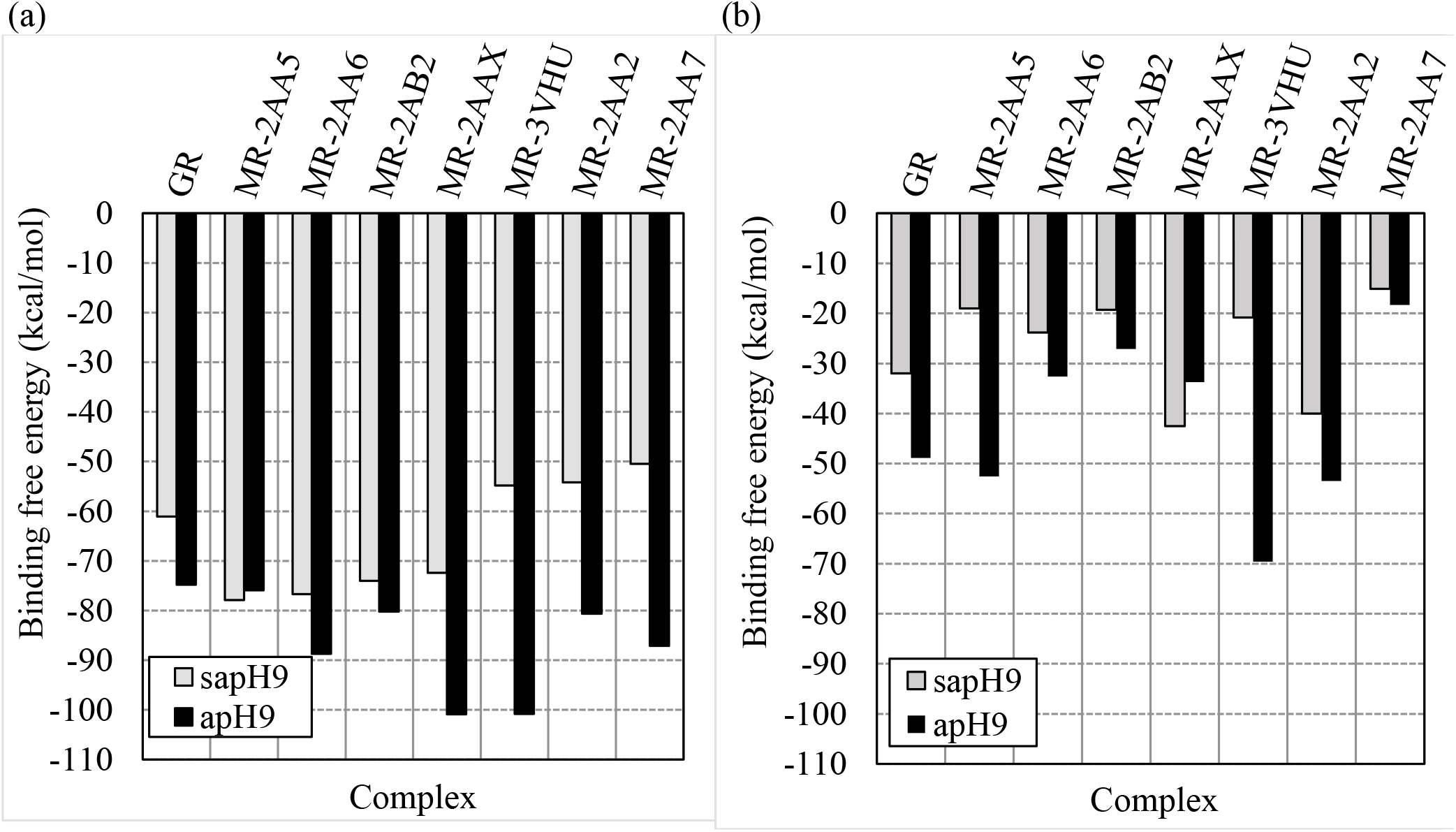
Stability of *apH9* and *sapH9* MR and GRα complexes using the MM/PBSA method applied to (a) minimized structures observed in crystals and (b) averages obtained from representative structures obtained from MD trajectories. The structures for MR are labelled according to Table 1. For GRα only two structural assemblies (based on PDB-ID:4P6W for *apH9*, and docked from MR PDB-ID:2AAX, see text for details) were analyzed for comparison.

**Figure 5:**
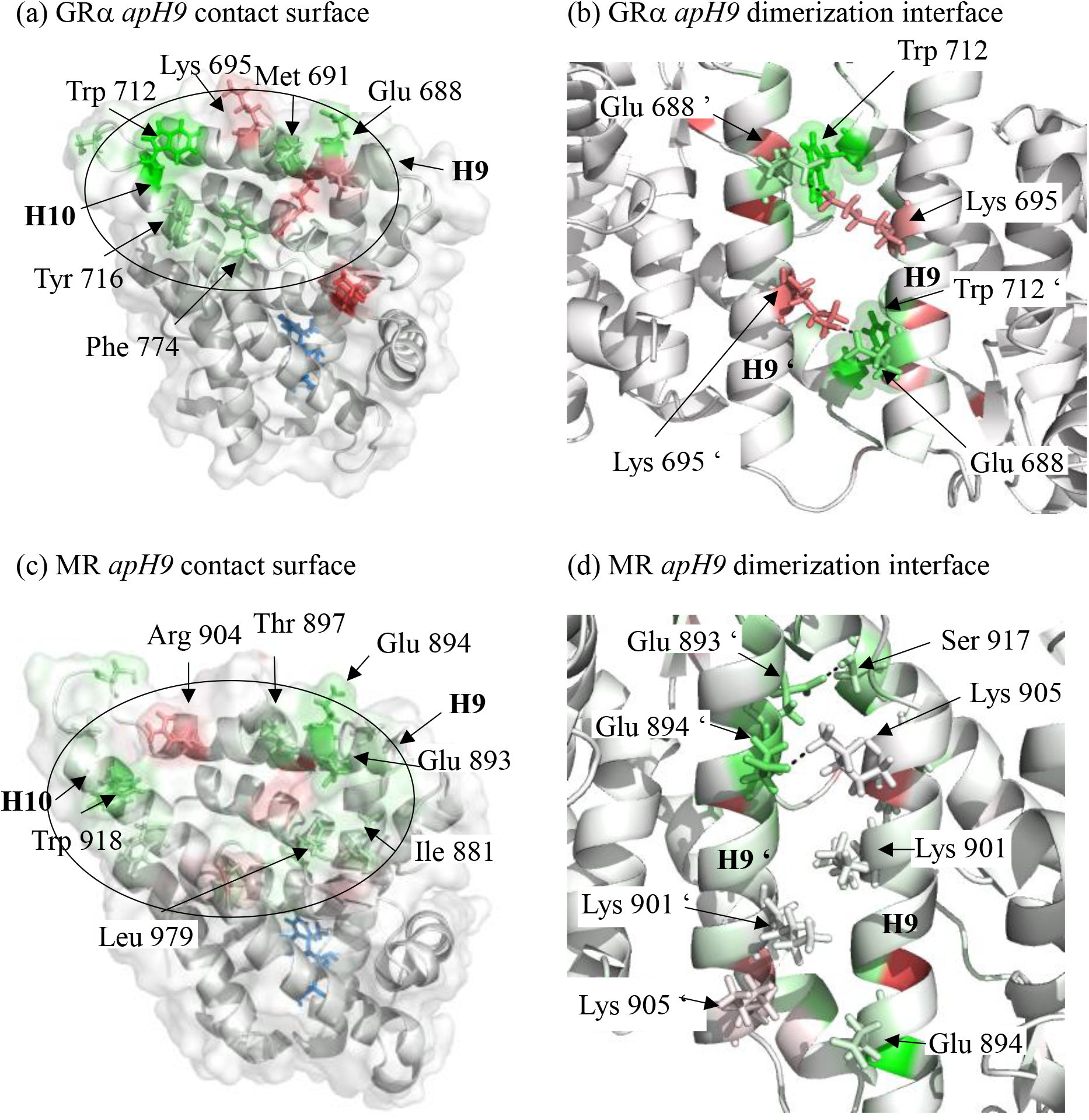
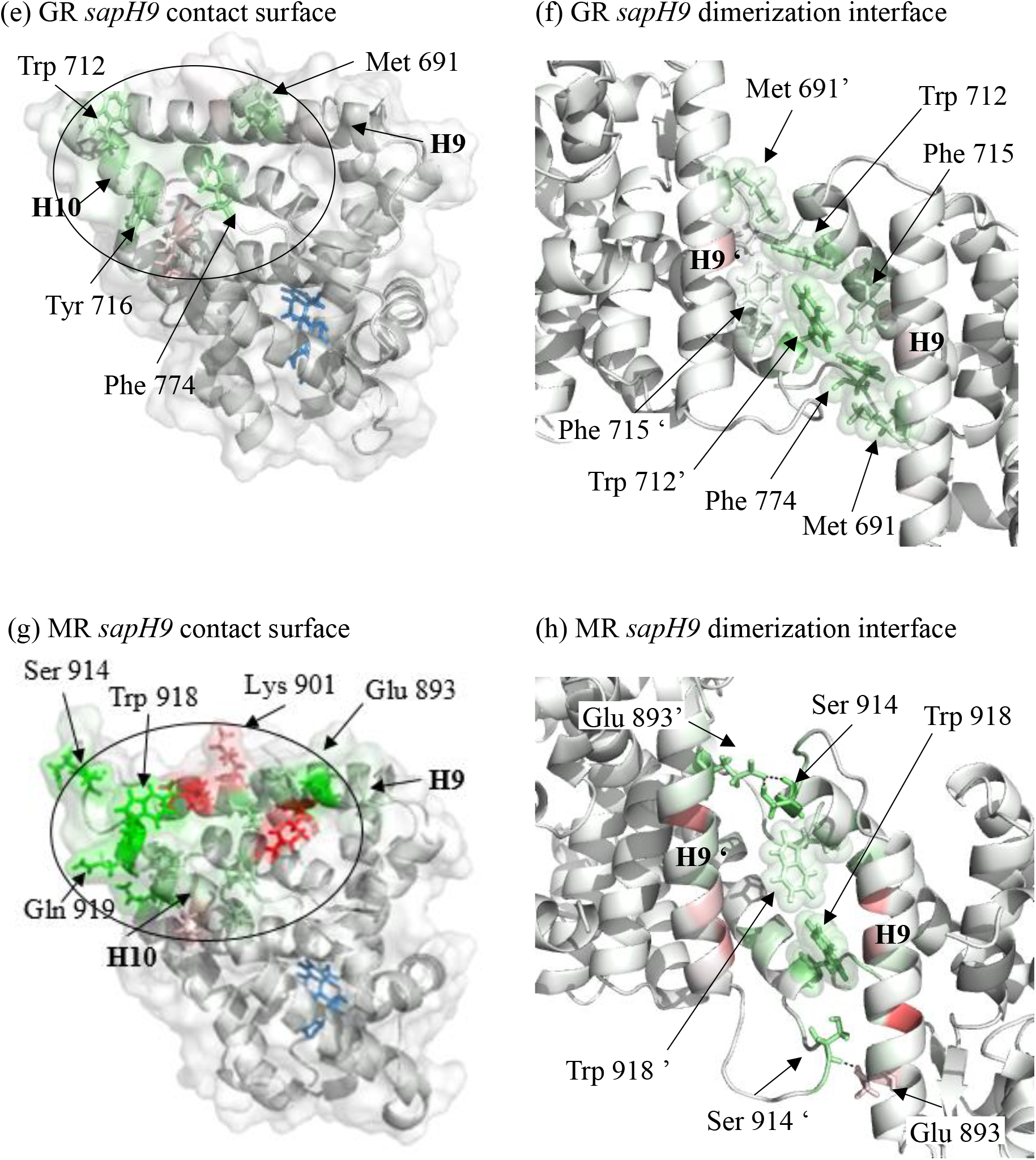
3D structure views of residues that contribute free energy to binding in *apH9* or *sapH9* assemblies. Contact surfaces are indicated on LBDs with an ellipse. Green and red colored amino-acids represent residues that stabilize or destabilize the dimer, respectively, according to a −10 kcal/mol to +10kcal/mol energy scale. Due to desolvation effect, one partner of a salt bridge may appear destabilizing but both partners together have a stabilizing effect. Dash lines show salt-bridges or polar contacts, (a) GRα *apH9* LBD contact surface, (b) GRα *apH9* dimerization interface, (c) MR *apH9* LBD contact surface, (d) MR *apH9* dimerization interface, (e) GRα *sapH9* LBD contact surface, (f) GRα *sapH9* dimerization interface, (g) MR *sapH9* LBD contact surface, (h) MR *sapH9* dimerization interface.

## 4 Discussion and Conclusion

In the super-family of NRs, the oxo-steroidian receptors, *i.e*. PR, AR, GRα and MR, constitute a group of 4 phylogenetically related protein sequences^1^. However, none of these receptors shows a LBD homodimer structure which is compliant with the NR canonical assembly, *i.e*. mediated by the H9, H10 and H11 interface^74,75^. Since oxosteroidian receptors are implicated in crucial transcriptional programs in both healthy and pathological situations, it is of major importance to determine which interface is responsible for LBD dimerization and how mutations may alter dimerization capabilities. In patients treated with long-term or high glucocorticoid doses, dysregulated GRα dimerization has been regarded as potentially responsible for side-effects^75^. In addition, inhibition of dimerization may also be a valuable approach for the design of drugs with less potential to develop glucocorticoid resistance in patients^75^. Therefore, fundamental knowledge on oxo-steroidian receptor dimerization is required to develop strategies to improve therapies. However, reconciling surface residue mutation information with dimerization interfaces is complicated because NRs are allosteric and residues that are not part of the dimerization interface might alter complex formation^27^. For GRα, a plethora of missense mutations that substitute residues positioned in the whole LBD have been reported to alter transactivation activity^76^ while transcriptional activation of target genes is effectively carried by receptor dimers. In addition, heterogeneous LBD assemblies that use alternative interfaces have been reported^77^. Therefore, there is currently no consensus for a shared homodimer assembly among the four oxo-steroidian receptors. However, there may be at least agreement on the fact that the oxo-steroidian C-terminus extremity, *i.e*. the F-domain, likely prevents the canonical homodimer from forming^37^ in GRα^27^, AR^28^ and PR^25^. No MR LBD homodimer structure has yet been reported.

In this report, we wished to analyze the stability of MR complex structures and compare them to its closest GRα homolog which has been extensively studied^26,27^. MR LBD crystals were collected from the PDB and analyzed with the PISA^50^, PRISM^51,52^ and EPPIC^53^ programs at both energetic and evolutionary levels. First, the canonical NR LBD homodimer was absent in all crystals. Second, in all MR LBD chains, the C-terminus extremity exhibited low flexibility and likely prevented the canonical homodimer from forming. Third, the homodimer assembly that was characterized experimentally for GRα^26^ was absent in all MR LBD crystals. On the contrary, a complex mediated by helices 9 in antiparallel orientations, (which we call *sapH9*), was frequently present in MR LBD crystals. This assembly presented structural similarities with a previously reported GRα assembly (which we called *apH9*^27^) although the distance between both H9 was greater in MR *sapH9* than in GRα *apH9*. In addition, a slight shift in helix turn between helices H9 of the two dimers was observed. Since MR and GRα are close homologues, MR LBD protein contacts in crystals might present similarities with those of GRα. The thorough examination of MR LBD crystals showed a heterogeneity of contacts between both receptors, with the exception of the *apH9/sapH9* mentioned above.

Using bioinformatics tools, the MR *sapH9* complex was examined to determine if it presented features of a biological protein-protein interaction, *i.e*. favorable free energy estimates (negative value) and high bioprobability (EPPIC) using evolutionary analysis. The *sapH9* was indeed the best ranked assembly among four different MR complexes observed in crystals. Of particular note, the conformation of the loop between H9 and H10 played a role in *sapH9* complex stabilization. In the conformation that was observed in a LBD bound to an antagonist ligand (PDB-ID:3WFF), the loop departed from the contact interface (“open” conformation) while in the conformation of PDB-ID:2AAX, *i.e*. a LBD bound to an agonist ligand, the loop was at the interface (“closed” conformation) and contributed significantly to the binding free energy of complex stabilization. Intriguingly, the *apH9* contact found for GRα was absent from all MR LBD crystals examined. Therefore, a model was built to determine if the MR *apH9* complex could be more stable than the experimentally observed *sapH9* and both MR *sapH9* and *apH9* assemblies were analyzed by the MM-PBSA method applied to energy minimized structures and MD simulations. From this analysis, the *apH9* contact showed significantly greater stability than the *sapH9*. This enhanced stability may be attributed to the formation of salt-bridges between Glu 894 and Lys 905 in the *apH9* complex but not in *sapH9*. Finally, both *apH9* and *sapH9* assemblies brought hydrophobic residues in contact at the interface, in particular the Trp 819, *i.e*. Trp 712 in GRα, which contributes significant binding free energy and which was previously shown to play a crucial role in GRα *apH9* complex stability. Thus, docked models of the MR *apH9* assembly showed greater stability than the *sapH9* using both energy minimized structures or MD trajectories. In addition, the biological likelihood of the former was significantly greater than the latter using EPPIC.

The reason why MR LBD crystals showed *sapH9* complexes but not the *apH9* assembly is intriguing. Two possibilities may be invoked. First, the usage of crystallization additives may result in the alteration of contact interfaces. Such additives were indeed retrieved from most MR LBD crystals (Supp. info S13). Several molecules of ethanediol were localized in close vicinity to the helices 9 and 10 and may have prevented the *apH9* assembly from forming. Second, LBD mutations designed to improve solubility and crystallization might be also responsible for assembly alterations, *e.g*. the frequent C910S mutant located in the loop between H9 and H10^78^. Finally, *sapH9* and *apH9* assemblies might represent two possible homodimeric states. The recurrent use of similar interaction interfaces in two close homologs that share DNA response elements^7^ while having specific regulatory function points to the need for further experimental investigation. In particular, while the recurrent use of a similar interface in GRα and MR points to its functional significance, it is not established whether this interface is used in the context of GRα or MR dimers or within larger assemblies where other partners provide additional stabilization. If a similar LBD dimerization interface is indeed shared by the two homologs, it could pave the way for providing for the first time a structural basis for the cross-talk between both receptors^33^.

## Acknowledgements

This work was supported by the French National Institute of Health and Medical Research (INSERM), the French National Center of Scientific Research (CNRS) and Strasbourg University (UdS). We are grateful to Dr. Roland H. Stote for thorough manuscript reading and corrections. We thank Steve Runser who wrote a script to carry out additive retrieval on LBDs. We would like also to acknowledge the High Performance Computing Center of the UdS and their staff for supporting this work by providing scientific support and access to computing resources. Part of the computing resources were funded by the Equipex Equip@Meso project (Programme Investissements d’Avenir). This study was supported by the grant ANR-10-LABX-0030-INRT, a French State fund managed by the Agence Nationale de la Recherche under the frame program Investissements d’Avenir ANR-10-IDEX-0002-02.

## Supplementary information

**S1 Figure:**
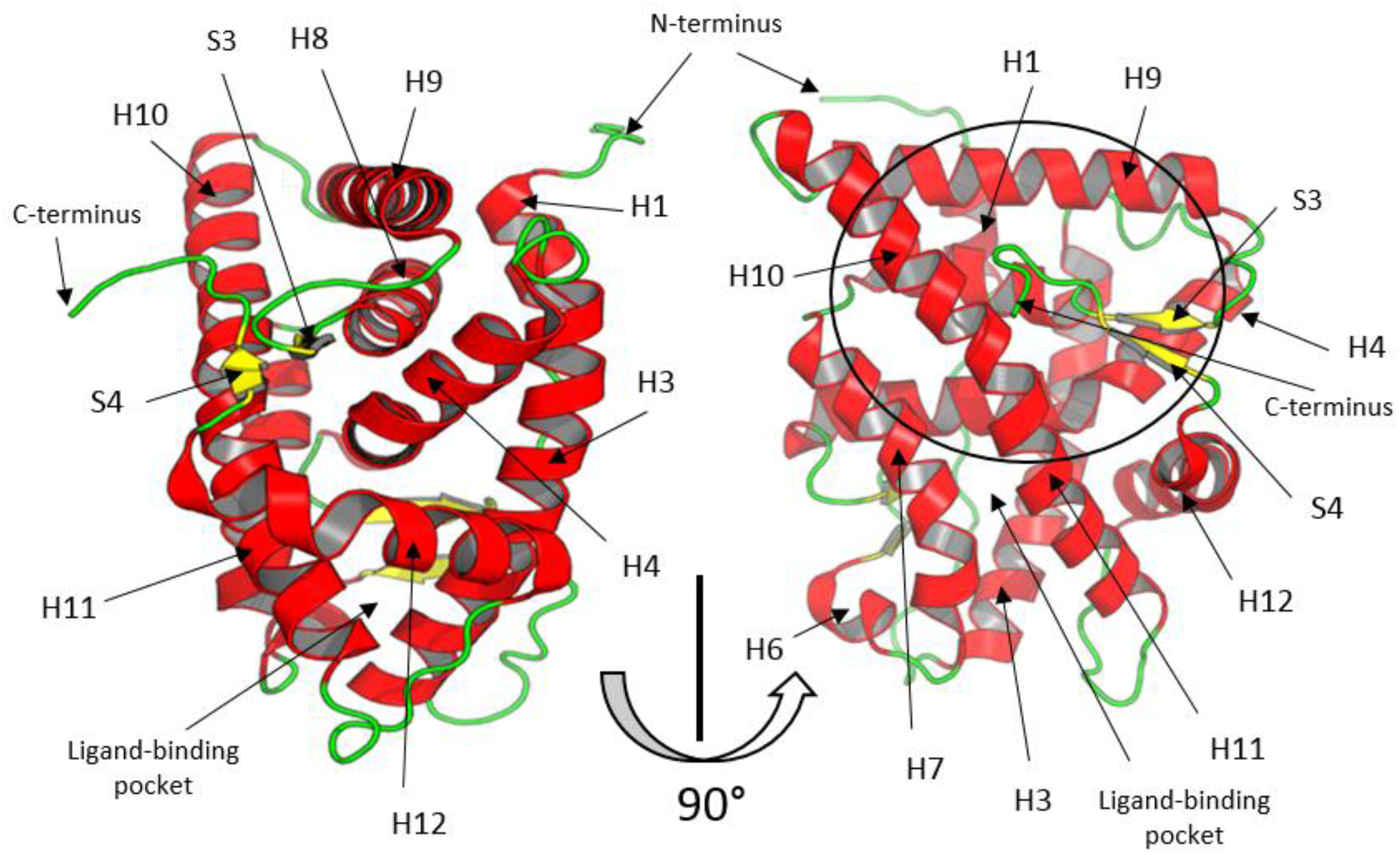
Schematic representation of GRα or MR LBD fold. There are 11 helices numbered from 1 to 12 – however, helix 2 does not exist in both receptors – and 4 β-strands (S1 to S4) that build 2 β-sheets. Helices, β-strands and loops are colored in red, yellow and green, respectively. The circle indicates the canonical NR homo- or heterodimerization contact surface which is mediated by H9, H10 and H11 residues which is observed in many NRs but not in GRα or MR. The F-domain that forms a steric hindrance to this canonical dimerization is Cter to H12 and contains S4.

**S2 Figure:**
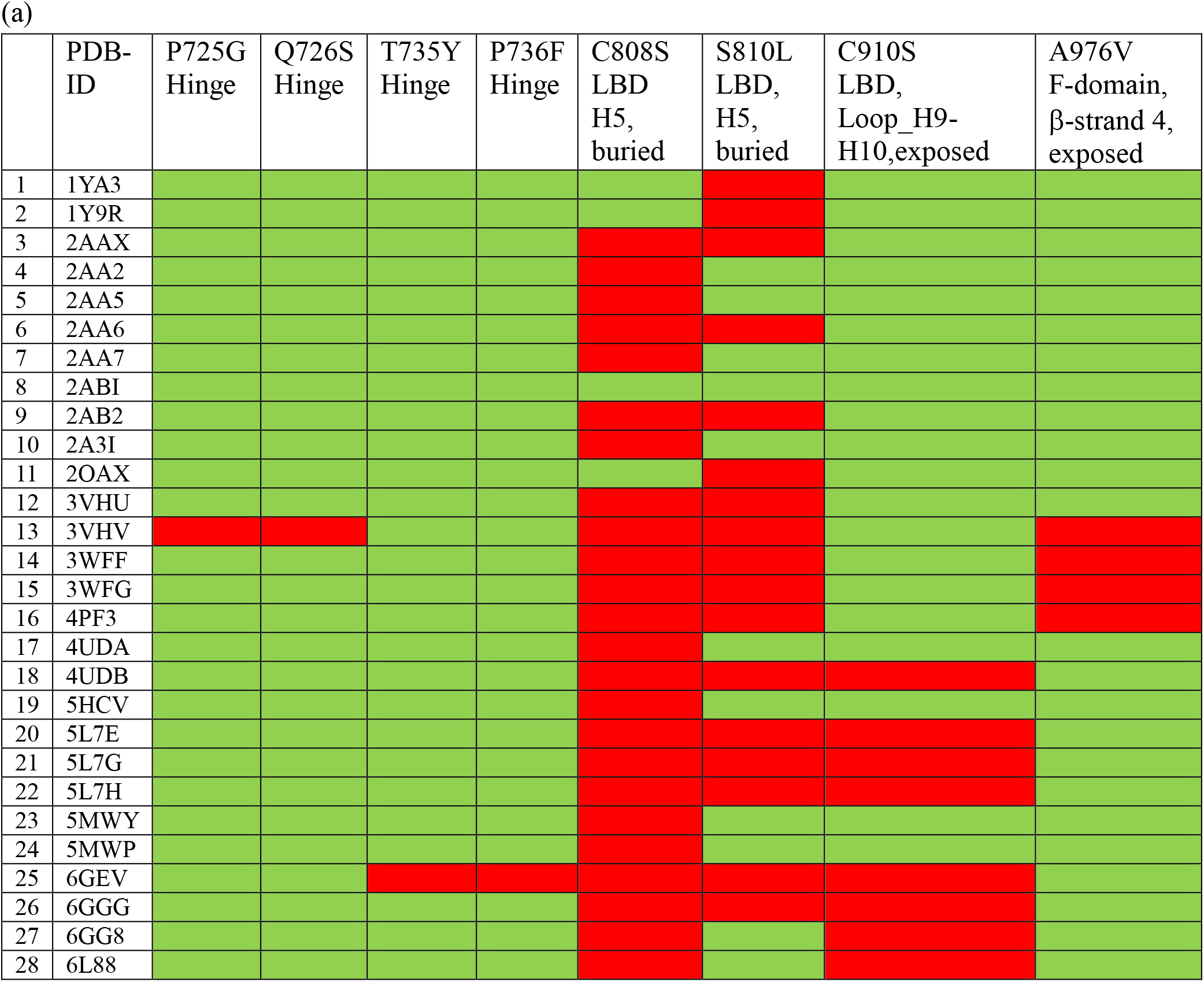
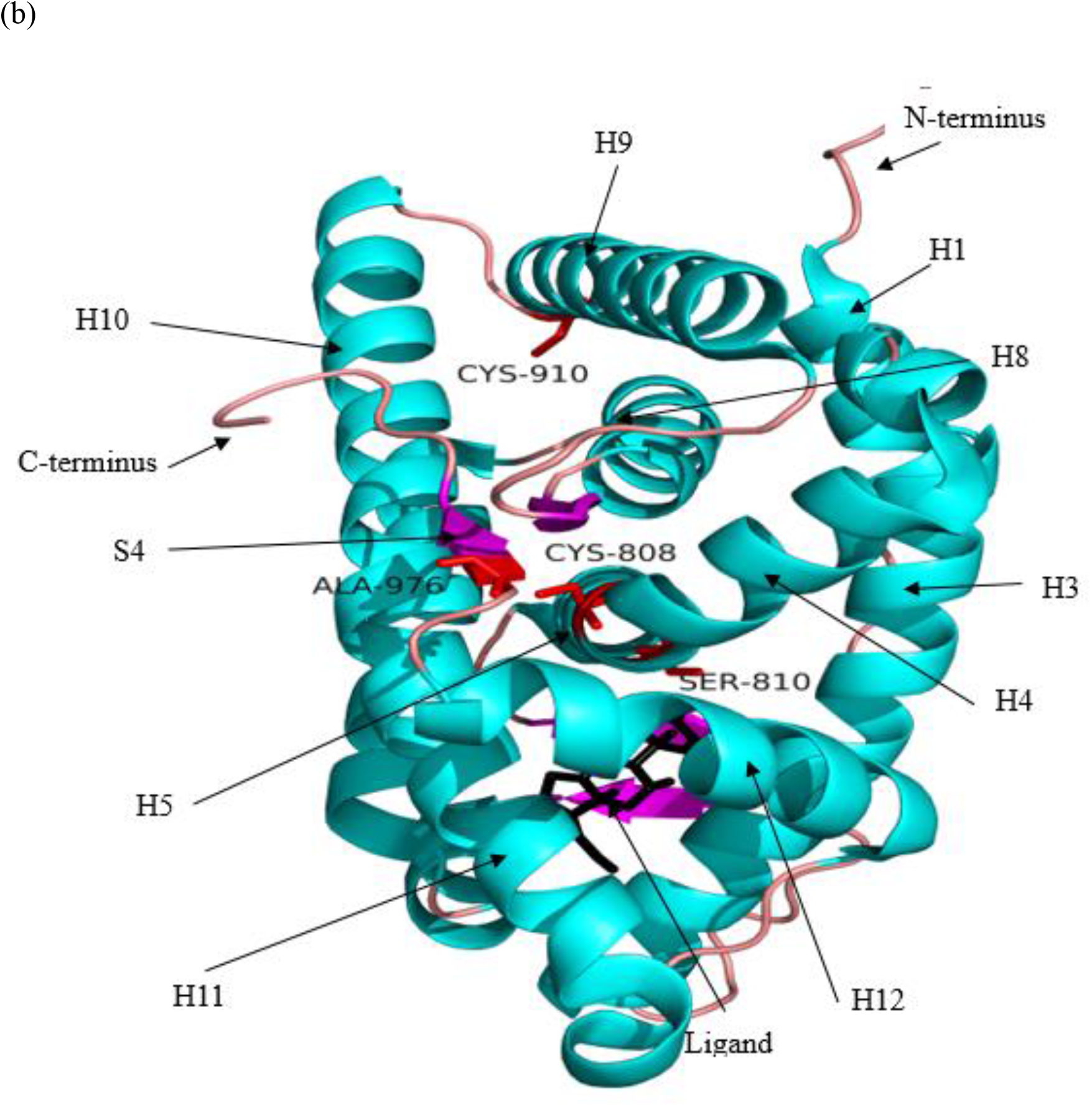
Mutated residues in crystallized MR LBD structures. (a) Mutation table: Green and red cells correspond to wild-type and mutated positions, respectively. Residue offsets refer to the human wildtype MR (RefSeq-ID:NP_000892). (b) Structural localization of mutated residues in MR LBD. Red colored residues represent mutated positions. Helices, β-strands, loops and ligands are colored in cyan, magenta, salmon and black, respectively.

**S3 Figure:**
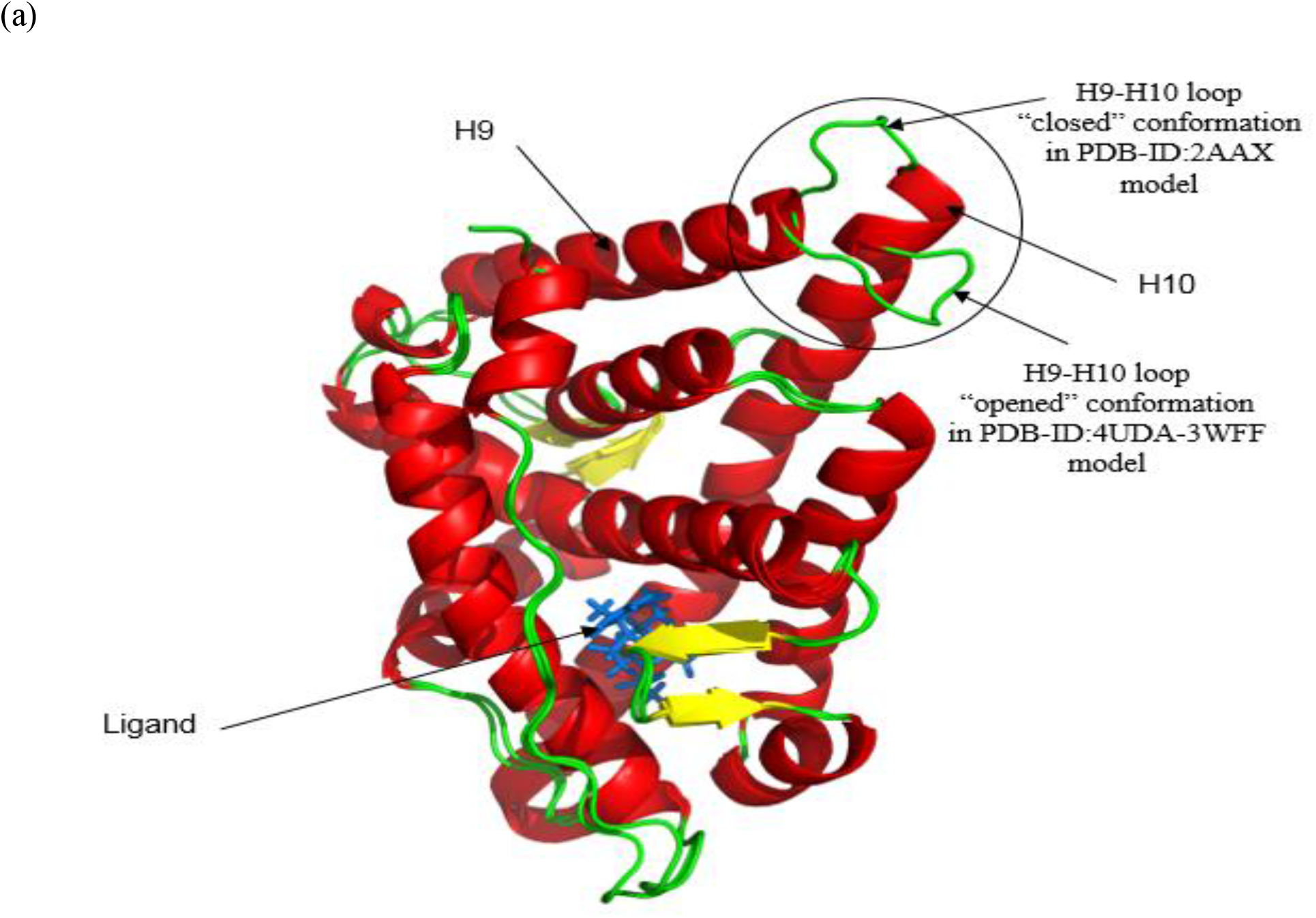
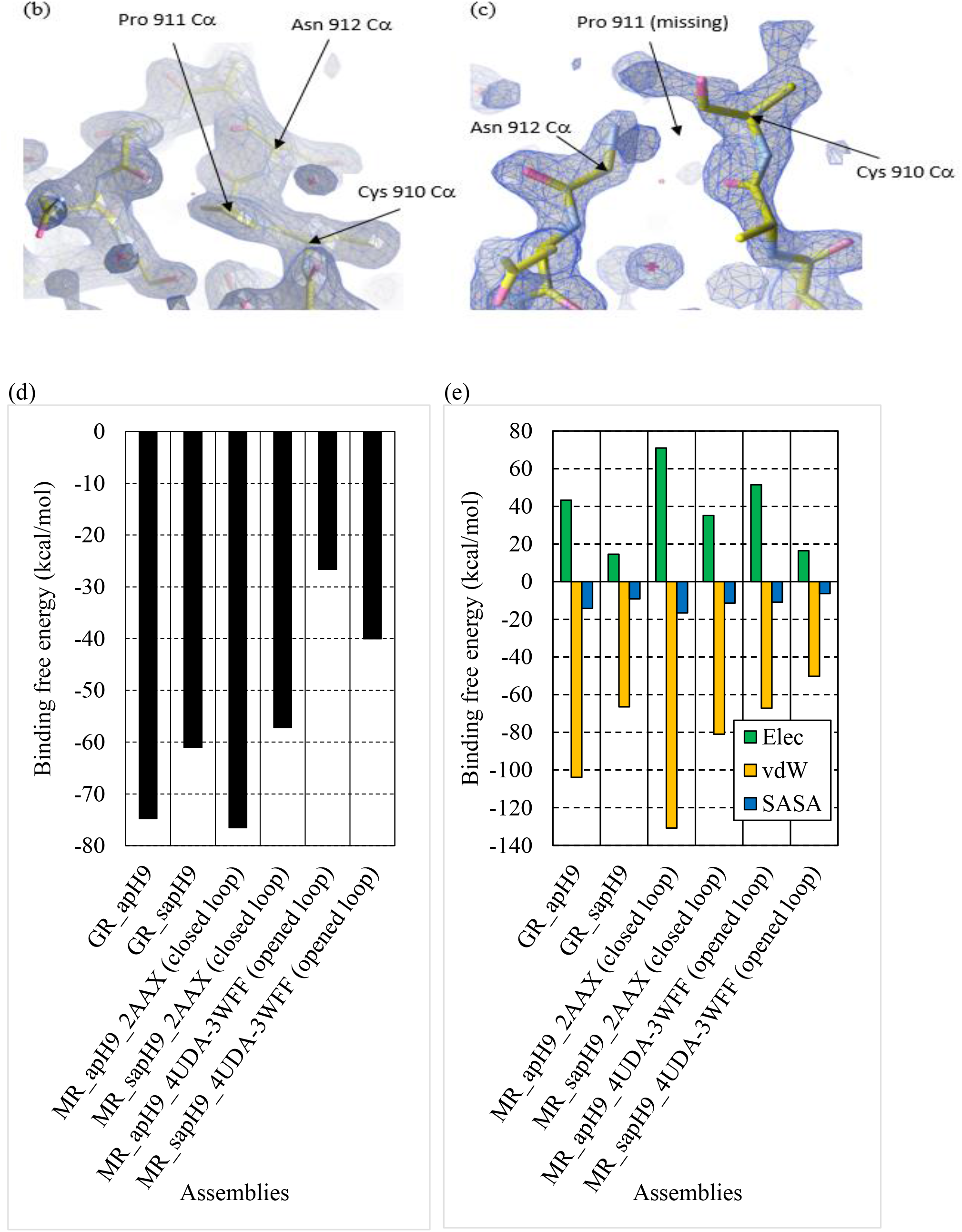
Alternative conformations of the loop between H9 and H10 in MR LBD as observed in crystals. (a) Superimposition of 4UDA-3WFF model (“opened” loop conformation) and PDB-ID:2AAX (“closed” loop conformation); The H9-H10 loop is surrounded by a circle. (b) Electronic densities of the loop in PDB-ID:3WFF and (c) PDB-ID:2AAX. (d) MM-PBSA total binding free energy in minimized MR *sapH9* and *apH9* complexes and (e) decomposition into electrostatic (Elec), van der Waals (vdW) and solvent accessible surface area (SASA) components. GRα was used as a reference.

**S4 Table:** Summary table of *sapH9* and *apH9* constructed models. The *apH9* assembly has been observed in only one crystal, *i.e*. GRα PDB-ID:4P6W, while the *sapH9* complex has been observed in seven MR crystals, *i.e*. PDB-ID:2AAX, 2AB2, 2AA2, 2AA5, 2AA6, 2AA7 and 3VHU. Wild-type protein LBDs were superposed onto homodimer templates to build the models. In total, 16 models were built.

|  | 3D molecular model | Assembly | Wild-type protein LBD | Homodimer template |
| --- | --- | --- | --- | --- |
| 1 | GR-4P6W-*apH9* | *apH9* | GRα | 4P6W |
| 2 | GR-4P6W-*sapH9* | *sapH9* | GRα | 2AAX |
| 3 | MR-2AAX-*sapH9* | *sapH9* | MR | 2AAX |
| 4 | MR-2AA7-*sapH9* | *sapH9* | MR | 2AA7 |
| 5 | MR-2AB2-*sapH9* | *sapH9* | MR | 2AB2 |
| 6 | MR-2AA6-*sapH9* | *sapH9* | MR | 2AA6 |
| 7 | MR-2AA2-*sapH9* | *sapH9* | MR | 2AA2 |
| 8 | MR-2AA5-*sapH9* | *sapH9* | MR | 2AA5 |
| 9 | MR-3VHU-*sapH9* | *sapH9* | MR | 3VHU |
| 10 | MR-2AAX-*apH9* | *apH9* | MR | 4P6W |
| 11 | MR-2AA7-*apH9* | *apH9* | MR | 4P6W |
| 12 | MR-2AB2-*apH9* | *apH9* | MR | 4P6W |
| 13 | MR-2AA6-*apH9* | *apH9* | MR | 4P6W |
| 14 | MR-2AA2-*apH9* | *apH9* | MR | 4P6W |
| 15 | MR-2AA5-*apH9* | *apH9* | MR | 4P6W |
| 16 | MR-3VHU-*apH9* | *apH9* | MR | 4P6W |

**S5 Figure:**
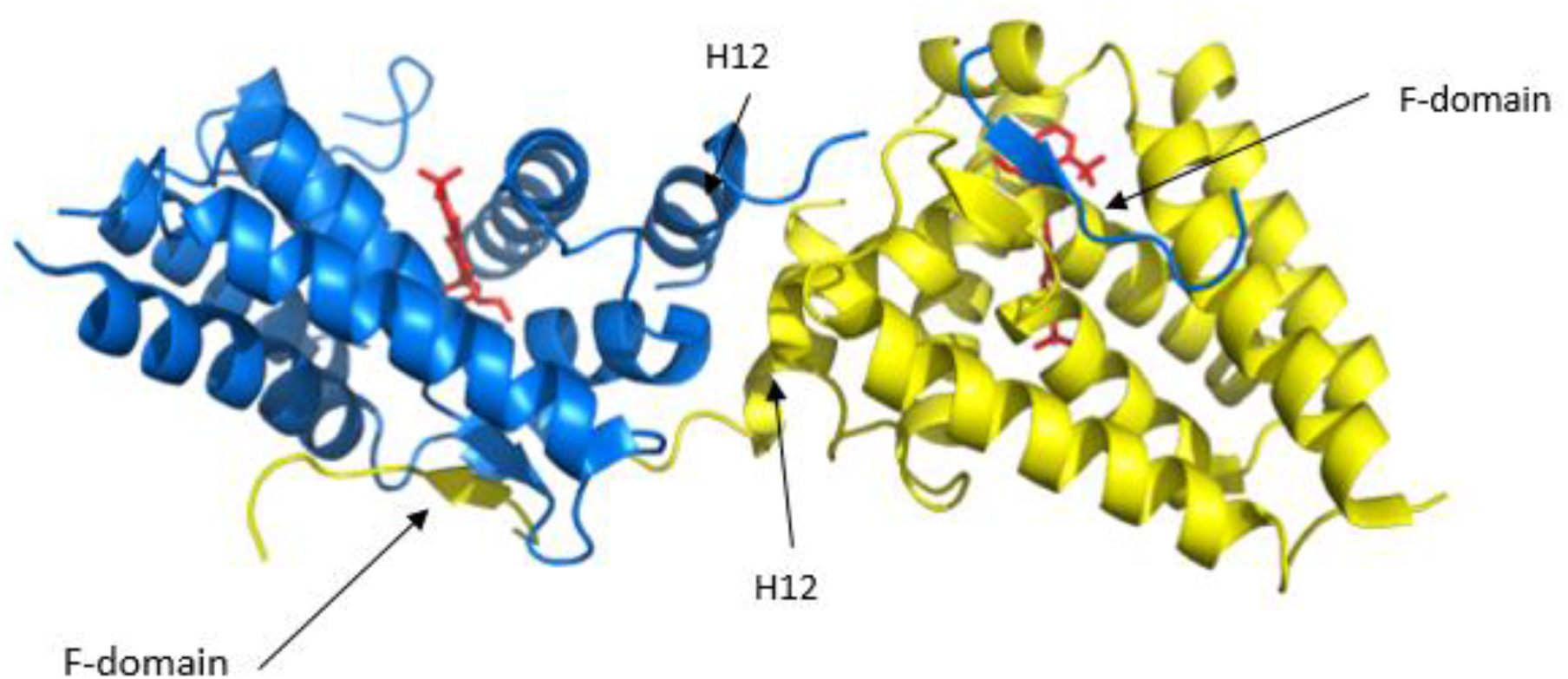
MR LBD swap dimer assembly. In crystal structure PDB-ID:6L88, both monomers swap their extreme C-terminus, *i.e*. the F-domain. Ligands are colored in red. Monomers are colored in blue and yellow.

**S6 Figure:**
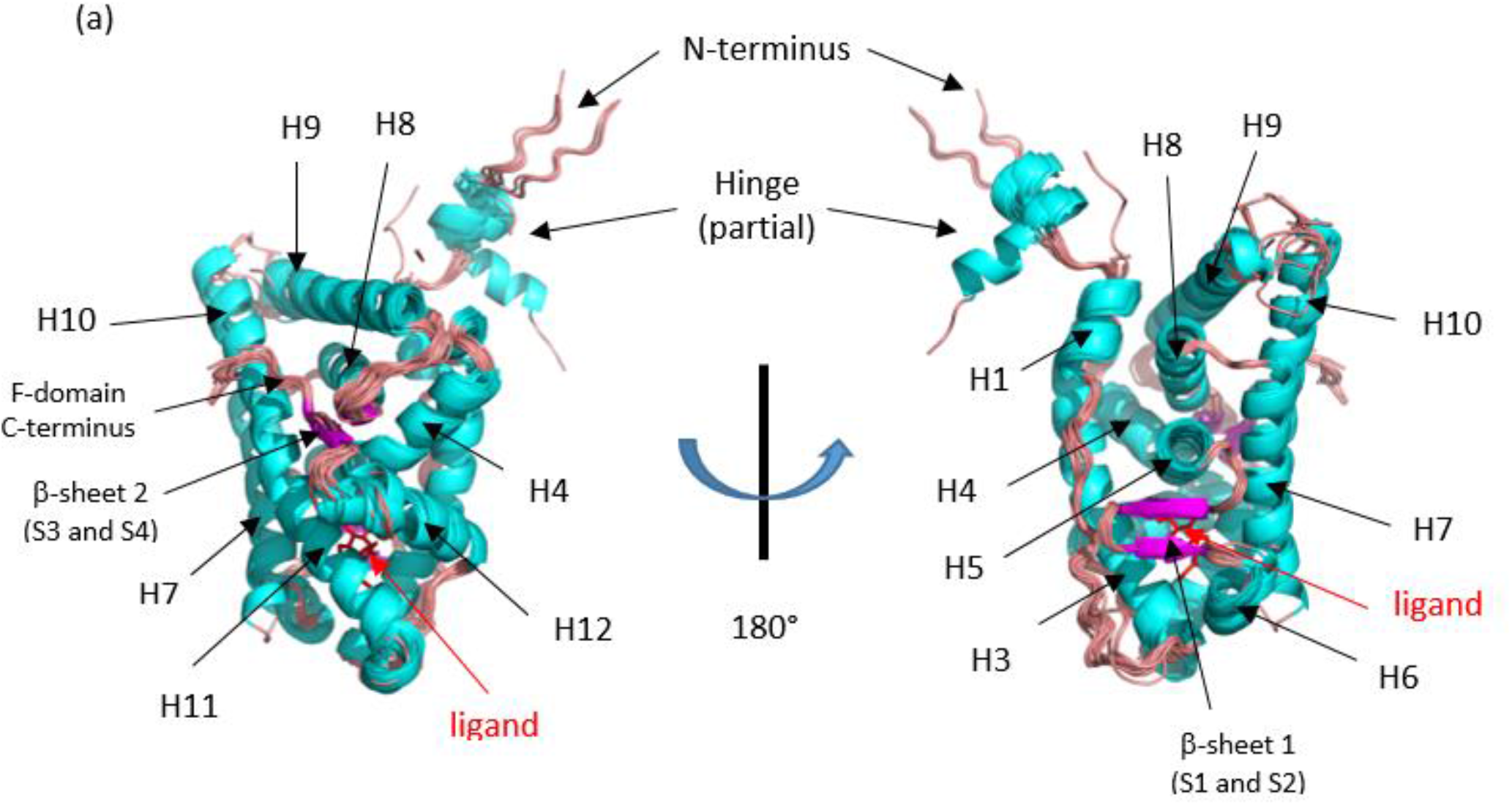
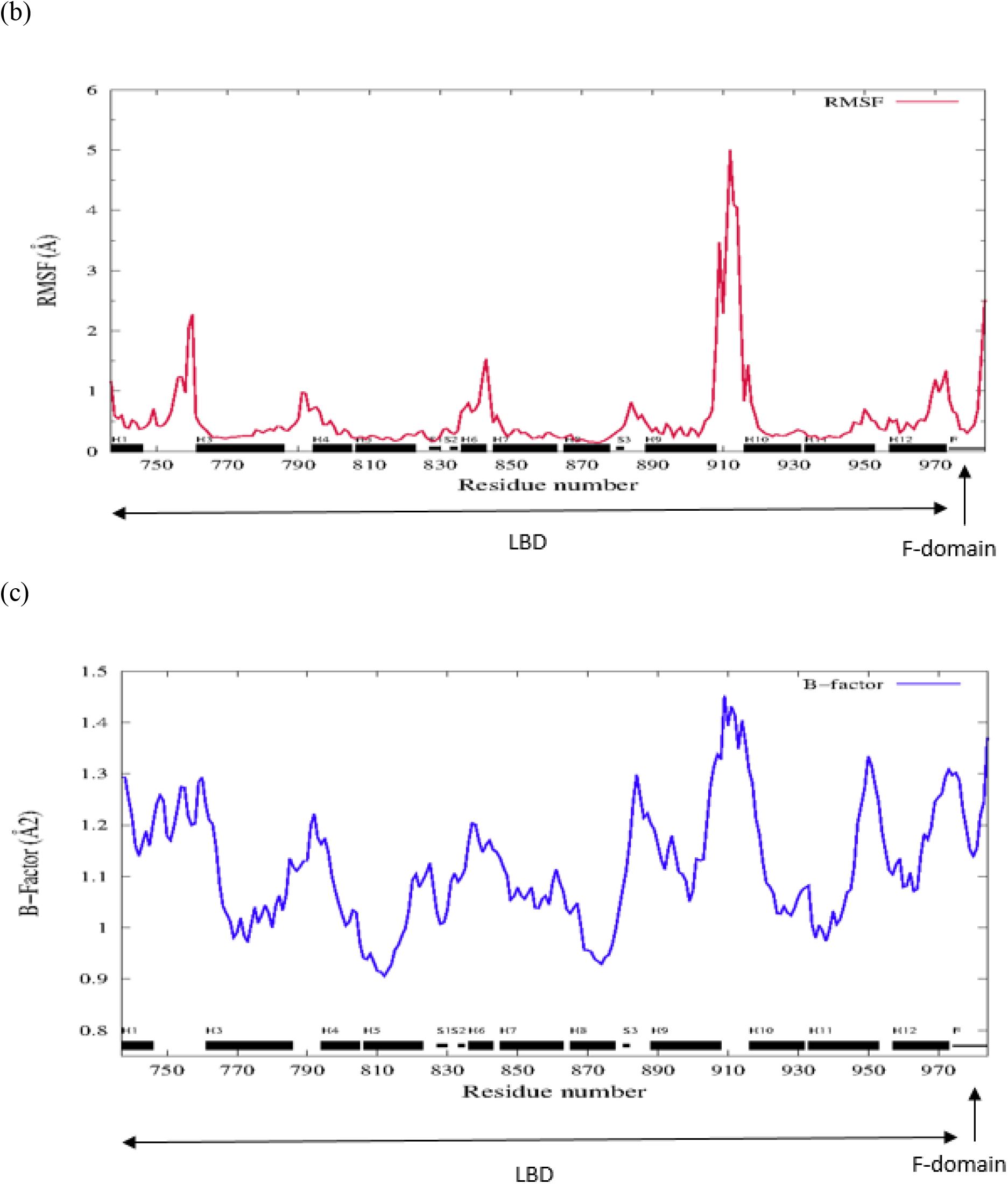
MR LBD structural statistics. (a) Superposition of LBD chains. α-helices, β-strands, loops and ligands are colored in cyan, magenta, salmon and red, respectively. The superposition was carried out by PSS^56^. In most receptor chains (84%), the loop between H9 and H10 was not fully resolved which emphasized its flexibility. However, it was fully resolved in PDB-ID:3WFF. At the C-terminus, both Arg 983 and Lys 984 were resolved in only 25.6 % of the chains, *e.g*. in PDB-ID:2A3I. All sequences were compared to the human wild-type MR (RefSeq-ID:NP_000892) to determine which residues had been mutated for expression and crystallization purposes. Most frequent mutations were C808S and S810L, *i.e*. two buried residues of H5, and C910S in the loop between H9 and H10 (Supp info S2). The C808S mutation, equivalent to F602S in human GRα (RefSeq-ID:NP_000167)^26^ has been reported to dramatically increase MR LBD expression^78^. (b) Root mean square fluctuation (RMSF) and (c) B-factor variations of Cα backbone along the full-length LBDs and F-domain. PDB-ID:6L88 was excluded from analysis.

**S7 Figure:**
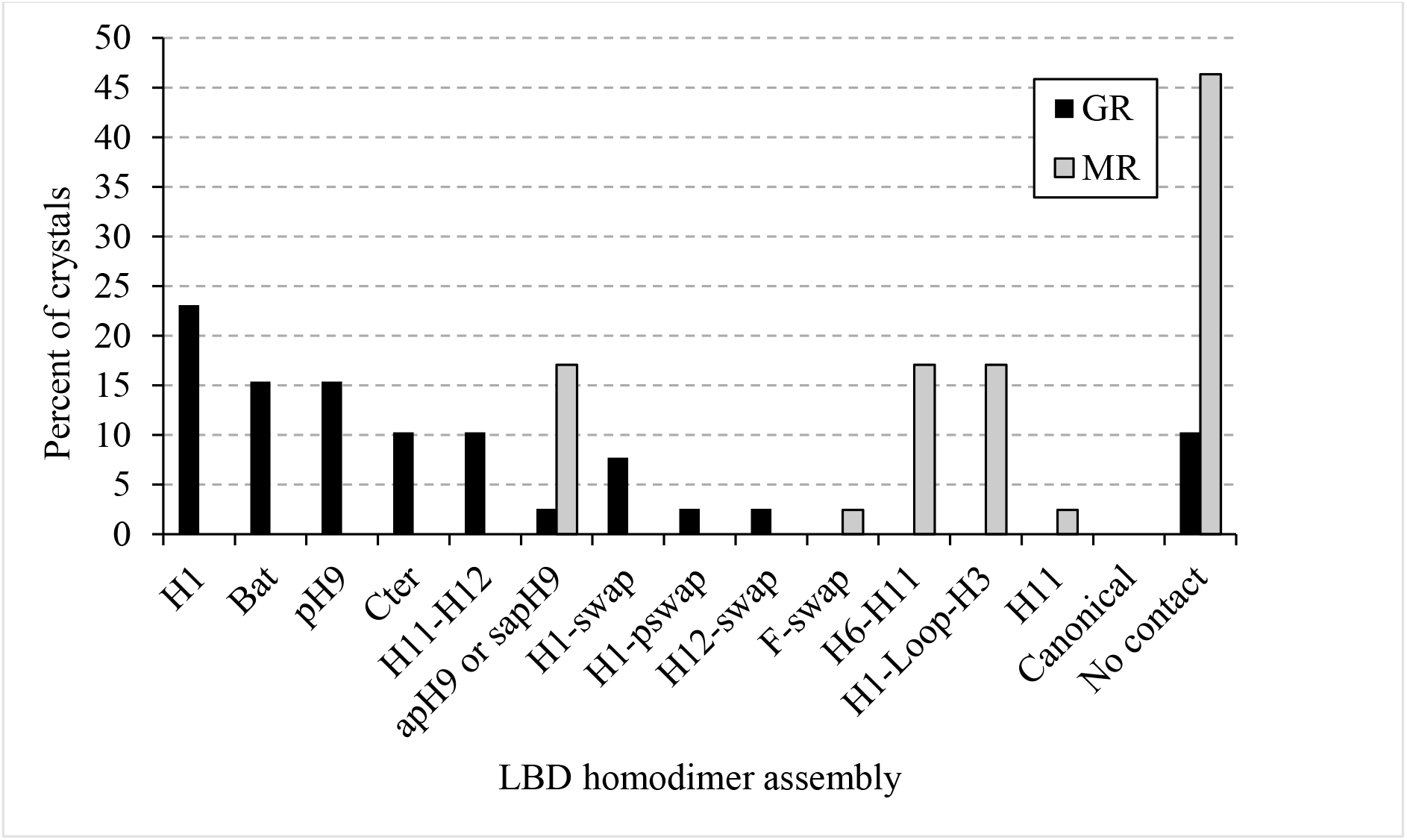
Frequencies of homodimeric assemblies in GRα and MR LBD crystals. 21 GRα crystals^27^ and 28 MR crystals were analyzed. Assemblies were named according to the secondary structures that were brought together at the contact interface.

**S8 Figure:**
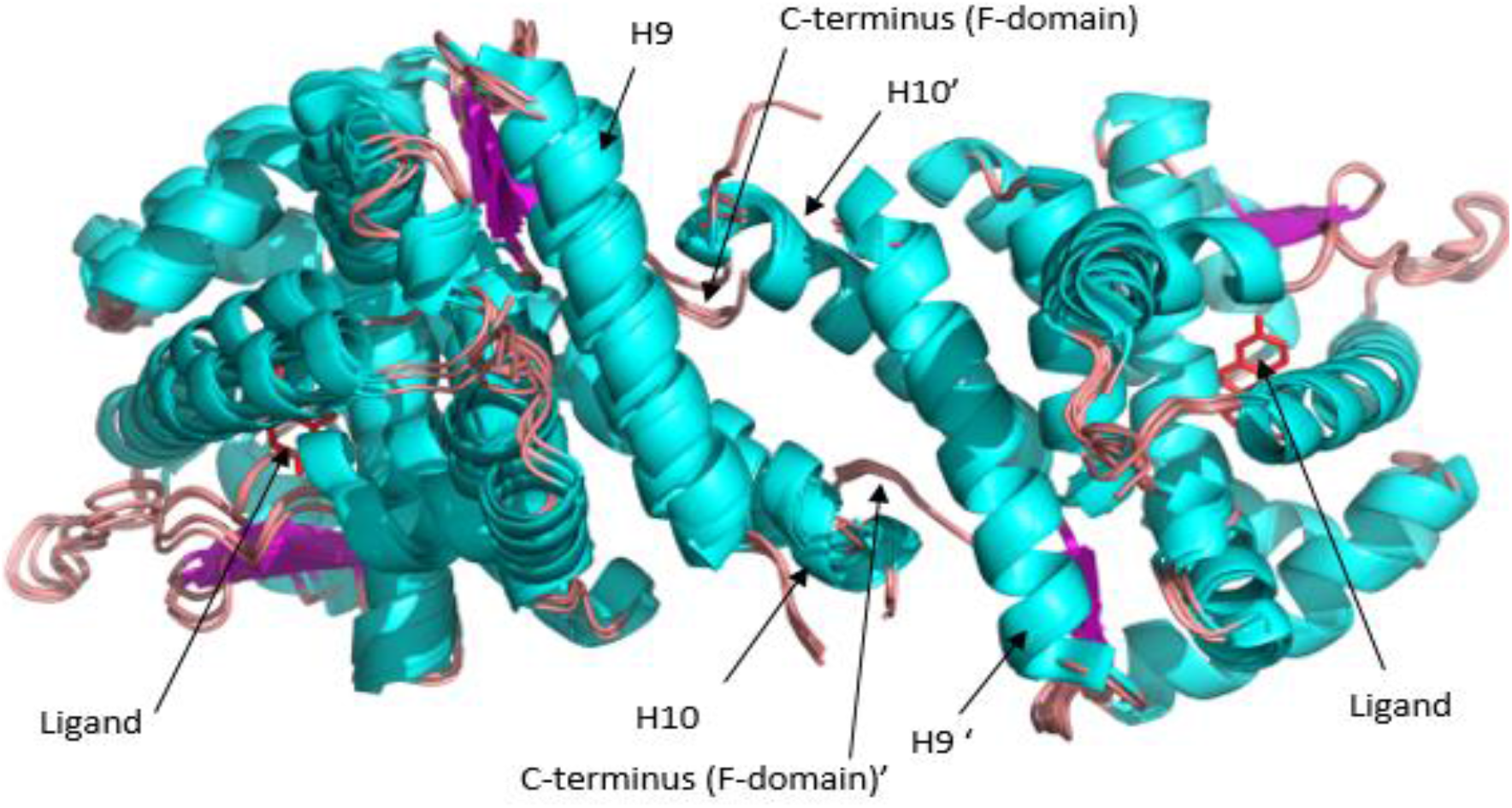
Structural superposition of all MR *sapH9* homodimers as observed in crystals. Helices, β-strands, loops and ligands are colored in cyan, magenta, salmon and red, respectively.

**S9 Table:** Binding free energy numerical data of MR LBD assemblies estimated by bioinformatics tools. a) PISA: the *sapH9* complex exhibited the largest (in absolute value) solvation free energy gain (−7.2 +/1.1 kcal/mol) while *H6-H11* (−3.5 +/-1.3 kcal/mol, p-value =1.6×10^-3^) and *H1-Loop-H3* (−0.9+/- 0.2 kcal/mol, p-value =10^-3^) assemblies were significantly less stable (One-sided Wilcoxon test), and b) PRISM: for each query and template pair, only max. and min. binding energy scores of dimers is reported (full data not shown) while average binding energy score is reported using all complexes (full data). The *H1* (−32.3+/-14.5 kcal/mol, p-value = 2.9×10^-2^) and *bat-like* (−24.4+/-6.5 kcal/mol, p-value = 4.89×10^-5^) assemblies were significantly less stable than the *sapH9* (One-sided Wilcoxon test).

| MR LBD query structure (PDB-ID) | Assembly in crystal | $\Delta iG$ (kcal/mol) | Average $\Delta iG$ (kcal/mol) and standard deviation across assemblies |
| --- | --- | --- | --- |
| 2AAX | sapH9 | -6.5 | -7.2 +/- 1.1 |
| 2AA2 |  | -7.7 |  |
| 2AA5 |  | -6.7 |  |
| 2AA6 |  | -7.4 |  |
| 2AA7 |  | -8.5 |  |
| 2AB2 |  | -8.5 |  |
| 3VHU |  | -5.4 |  |
| 2AAX | H6-H11 | -2.4 | -3.5 +/- 1.3 |
| 2AA2 |  | -2.1 |  |
| 2AA5 |  | -3.2 |  |
| 2AA6 |  | -3.4 |  |
| 2AA7 |  | -6 |  |
| 2AB2 |  | -3.2 |  |
| 3VHU |  | -4.2 |  |
| 2AAX | H1-Loop-H3 | -0.8 | -0.9 +/- 0.2 |
| 2AA2 |  | -0.9 |  |
| 2AA5 |  | -0.8 |  |
| 2AA6 |  | -0.9 |  |
| 2AA7 |  | -1.1 |  |
| 2AB2 |  | -0.9 |  |
| 3VHU |  | -0.6 |  |
| 3WFF | H11 | -4.4 | -4.4 +/- 0 |

(S9 Table continues on next page) (b)
| MR LBD query struct. (PDB-ID) | Query polypep. chain 1 | Query polypep. chain 2 | Template dimer PDB-ID and polypep. chains | Template dimer assembly and protein | Binding free energy (kcal/mol) | Average binding free energy (kcal/mol) |
| --- | --- | --- | --- | --- | --- | --- |
| 1YA3 | B | A | 2AA5:A,B | **sapH9 (MR)** | -12.8 | -41.2<br>+/- 15.6 |
| 2A3I | A | A | 2AB2:A,B |  | -30.5 |  |
| 2AA6 | B | A | 2AB2:A,B |  | -46.2 |  |
| 2AA6 | A | B | 2AB2:A,B |  | -40.4 |  |
| 2AA6 | A | B | 2AA5:A,B |  | -43.7 |  |
| 2AAX | B | A | 2AB2:A,B |  | -57.2 |  |
| 2AAX | A | B | 2AB2:A,B |  | -56.5 |  |
| 2AAX | A | B | 2AA5:A,B |  | -56.3 |  |
| 2AAX | B | A | 2AA5:A,B |  | -55.6 |  |
| 2AB2 | A | B | 2AA5:A,B |  | -45.4 |  |
| 2AB2 | B | A | 2AB2:A,B |  | -42.7 |  |
| 3VHV | A | A | 2AB2:A,B |  | -9.4 |  |
| 5L7E | A | A | 2AB2:A,B |  | -39.1 |  |
| 2AA7 | A | A | 1XV9:A,C | **H11 (CAR/RXRα)** | -8.5 | -6.1<br>+/- 3.5 |
| 2AB2 | A | B | 1XV9:A,C |  | -3.6 |  |
| 3WFF | A | A | 1ZUC:A,B | **H11-H12 (PR)** | -5.7 | -5.7 |
| 1Y9R | B | A | 1P93:A,C | **Bat-like (GR)** | -21.8 | -24.4<br>+/-6.5 |
| 1Y9R | A | B | 1P93:A,C |  | -8.47 |  |
| 2A3I | A | A | 1P93:A,C |  | -24.7 |  |
| 2A3I | A | A | 1P93:A,C |  | -20.6 |  |
| 2AA2 | A | A | 1P93:A,C |  | -28.3 |  |
| 2AA2 | A | A | 1P93:A,C |  | -24.2 |  |
| 2AA5 | A | B | 1P93:A,C |  | -32.4 |  |
| 2AA5 | B | A | 1P93:A,C |  | -21.8 |  |
| 2AA6 | A | B | 1P93:A,C |  | -33.8 |  |
| 2AA6 | B | A | 1P93:A,C |  | -16.9 |  |
| 2AA7 | A | A | 1P93:A,C |  | -29.7 |  |
| 2AA7 | A | A | 1P93:A,C |  | -26.5 |  |
| 2AAX | B | A | 1P93:A,C |  | -25 |  |
| 2AAX | B | A | 1P93:A,C |  | -20.3 |  |
| 2AB2 | A | B | 1P93:A,C |  | -33.9 |  |
| 2AB2 | B | A | 1P93:A,C |  | -20.5 |  |
| 2OAX | A | B | 1P93:A,C |  | -28.4 |  |
| 2OAX | A | B | 1P93:A,C |  | -15.7 |  |
| 3VHU | A | A | 1P93:A,C |  | -28.3 |  |
| 3VHU | A | A | 1P93:A,C |  | -22.1 |  |
| 3VHV | A | A | 1P93:A,C |  | -23.6 |  |
| 3WFF | A | A | 1P93:A,C |  | -38.1 |  |
| 3WFF | A | A | 1P93:A,C |  | -21.5 |  |
| 3WFG | A | A | 1P93:A,C |  | -23.6 |  |
| 3WFG | A | A | 1P93:A,C |  | -22 |  |
| 4PF3 | A | A | 1P93:A,C |  | -18.2 |  |
| 4PF3 | A | A | 1P93:A,C |  | -4.2 |  |
| 4UDA | A | A | 1P93:A,C |  | -26.5 |  |
| 4UDA | A | A | 1P93:A,C |  | -16 |  |
| 4UDB | A | A | 1P93:A,C |  | -15.3 |  |
| 5HCV | A | B | 1P93:A,C |  | -35.7 |  |
| 5HCV | B | A | 1P93:A,C |  | -35.4 |  |
| 5L7G | A | A | 1P93:A,C |  | -37,5 |  |
| 5L7G | A | A | 1P93:A,C |  | -18 |  |
| 5L7E | A | A | 1P93:A,C |  | -35.1 |  |
| 5L7E | A | A | 1P93:A,C |  | -24.7 |  |
| 5L7H | A | A | 1P93:A,C |  | -36.5 |  |
| 5L7H | A | A | 1P93:A,C |  | -22 |  |
| 2A3I | A | A | 3CLD:A,B | **H1**<br>(GR) | -5.9 | -32.3<br>+/-14.5 |
| 2AA2 | A | A | 3CLD:A,B |  | -35,2 |  |
| 2AA2 | A | A | 3E7C:A,B |  | -23.5 |  |
| 2AA5 | B | B | 3E7C:A,B |  | -55 |  |
| 2AA5 | A | B | 3CLD:A,B |  | -20.7 |  |
| 2AA6 | A | B | 3E7C:A,B |  | -57.3 |  |
| 2AA6 | B | A | 3CLD:A,B |  | -15.6 |  |
| 2AA7 | A | A | 3E7C:A,B |  | -50.7 |  |
| 2AA7 | A | A | 3CLD:A,B |  | -49.8 |  |
| 2AAX | A | B | 3E7C:A,B |  | -40.7 |  |
| 2AAX | A | B | 3E7C:A,B |  | -17.8 |  |
| 2AB2 | A | B | 3CLD:A,B |  | -38.6 |  |
| 2AB2 | A | B | 3E7C:A,B |  | -14.7 |  |
| 3VHU | A | A | 3CLD:A,B |  | -38.6 |  |
| 3VHU | A | A | 3E7C:A,B |  | -20.6 |  |
| 3WFF | A | A | 3CLD:A,B |  | -10.1 |  |
| 3WFF | A | A | 3E7C:A,B |  | -8.1 |  |

**S10 Figure:**
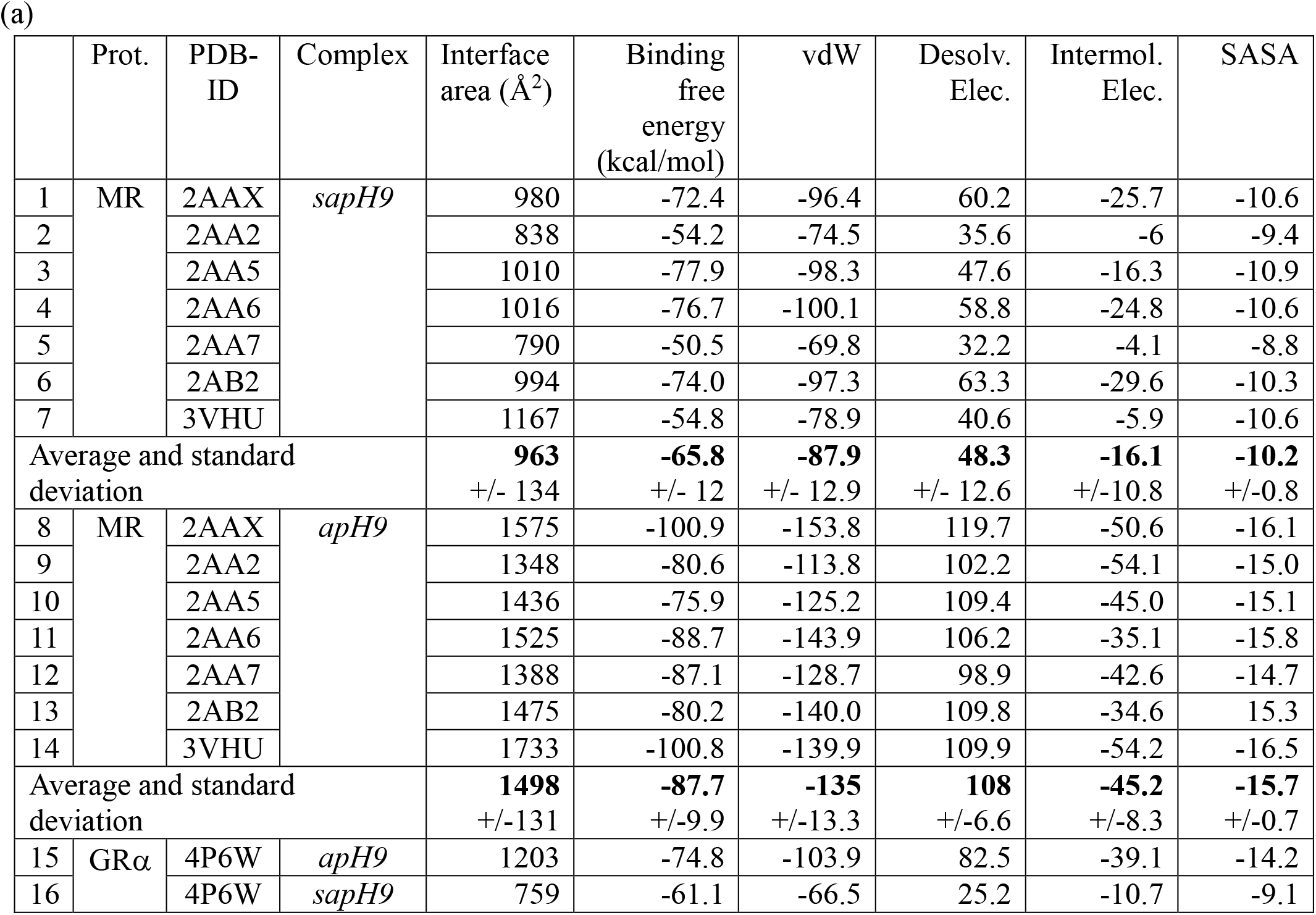
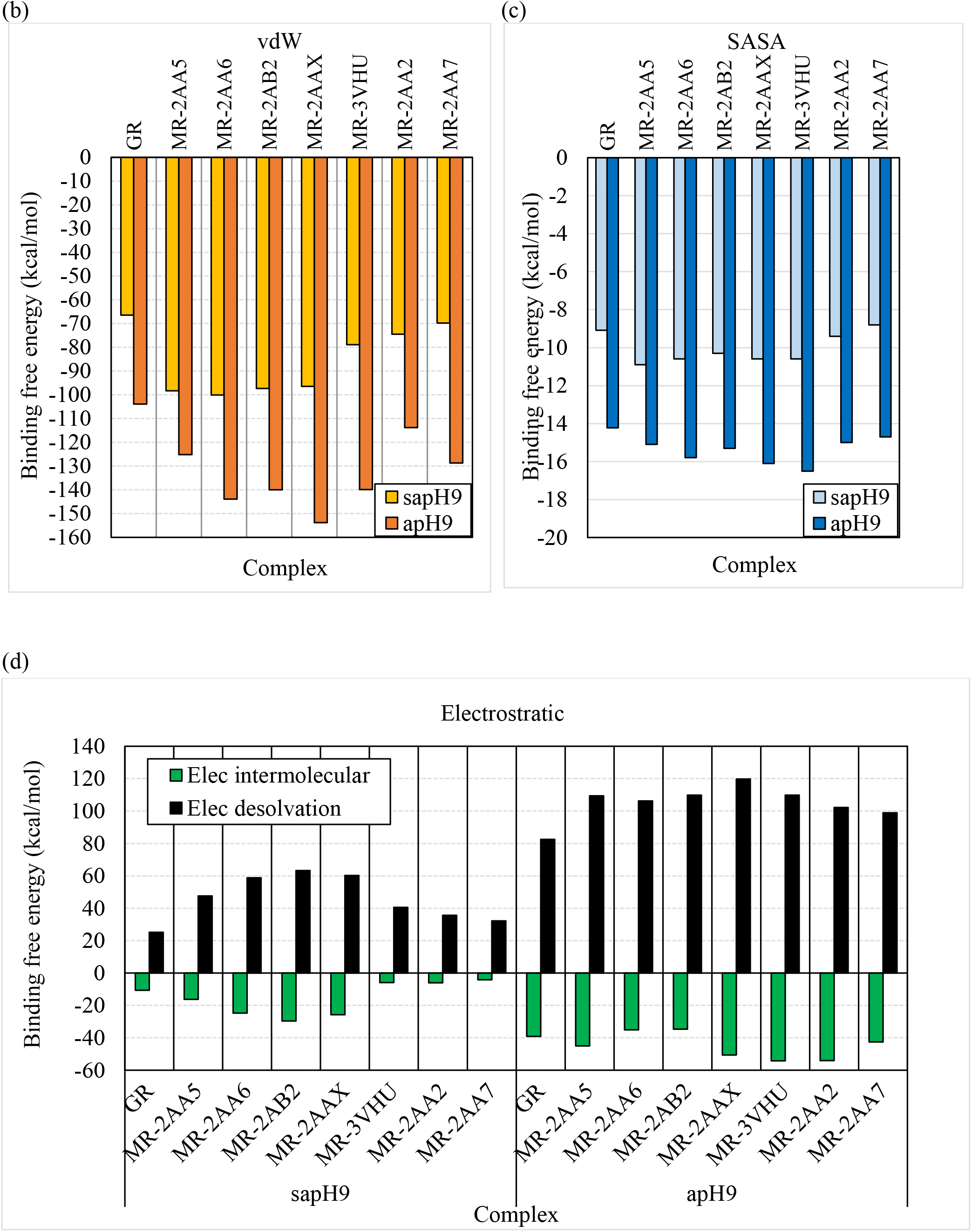
Stability of *apH9* and *sapH9* complexes using the MM-PBSA method applied to minimized structures observed in crystals and decomposition of binding free energy into its component, *i.e*. van der Waals (vdW), Electrostatic (Desolvation and intermolecular) and Solvent accessible surface area (SASA). (a) Numerical data. Graphical representation of binding free energy terms: (b) vdW, (c) SASA and (d) Electrostatic.

**S11 Figure:**
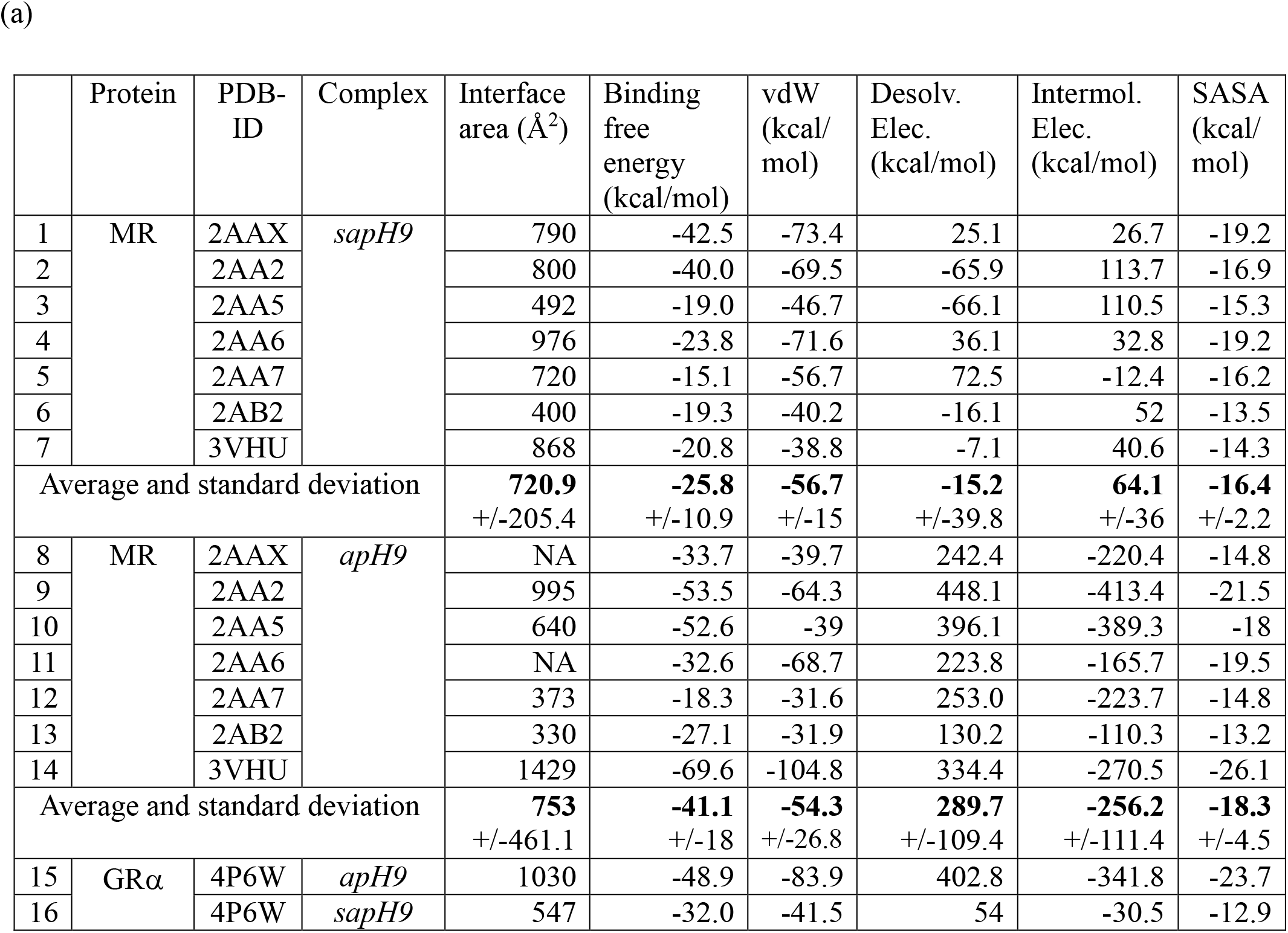
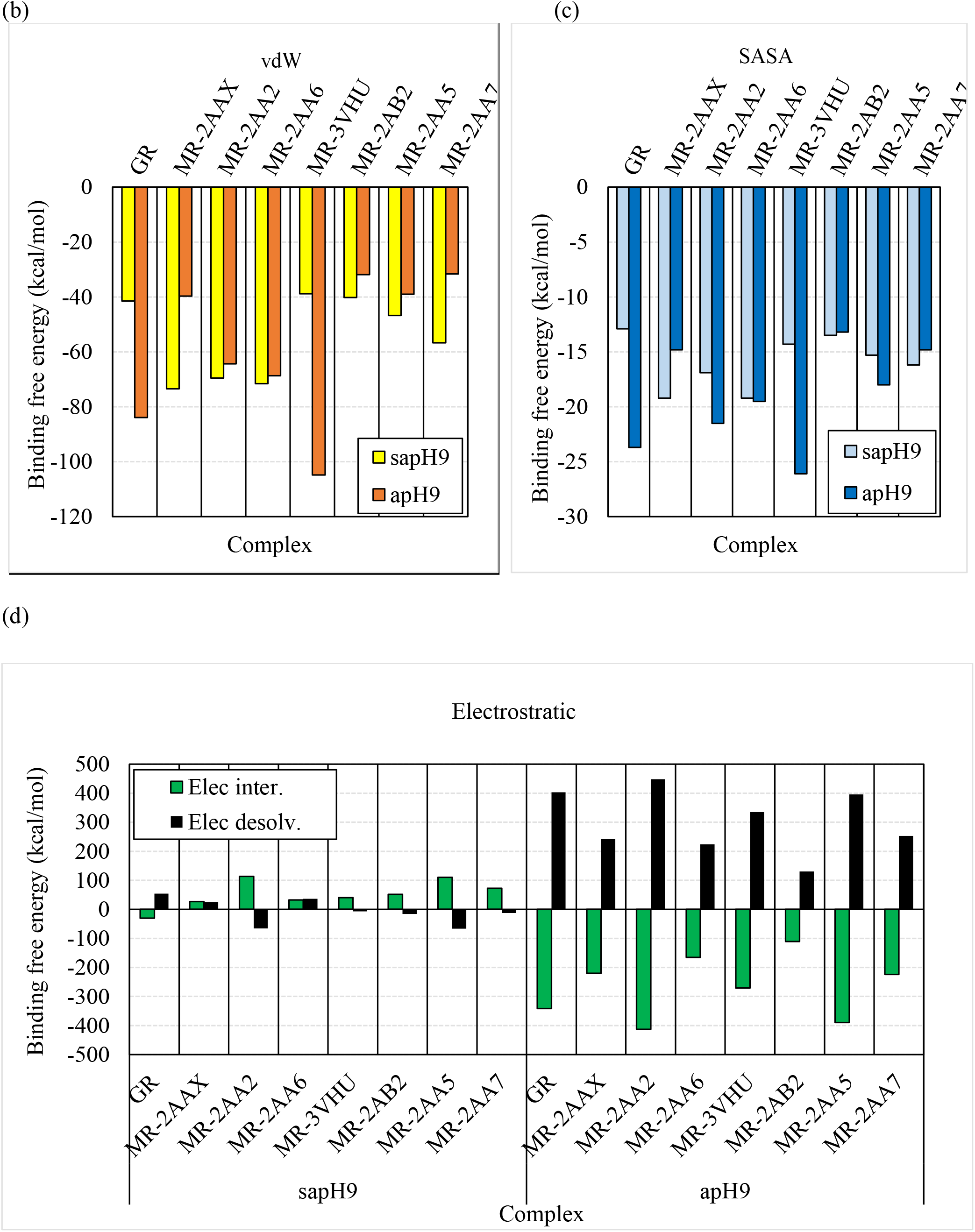
Stability of *apH9* and *sapH9* complexes using the MM/PBSA method applied to average structures obtained from a MD trajectory and decomposition of binding free energy into its component, *i.e*. van der Waals (vdW), Electrostatic (Desolvation and intermolecular) and Solvent sccessible surface area (SASA). (a) Numerical data. Graphical representation of binding free energy terms: (b) vdW, (c) SASA and (d) Electrostatic.

**S12 Figure:**
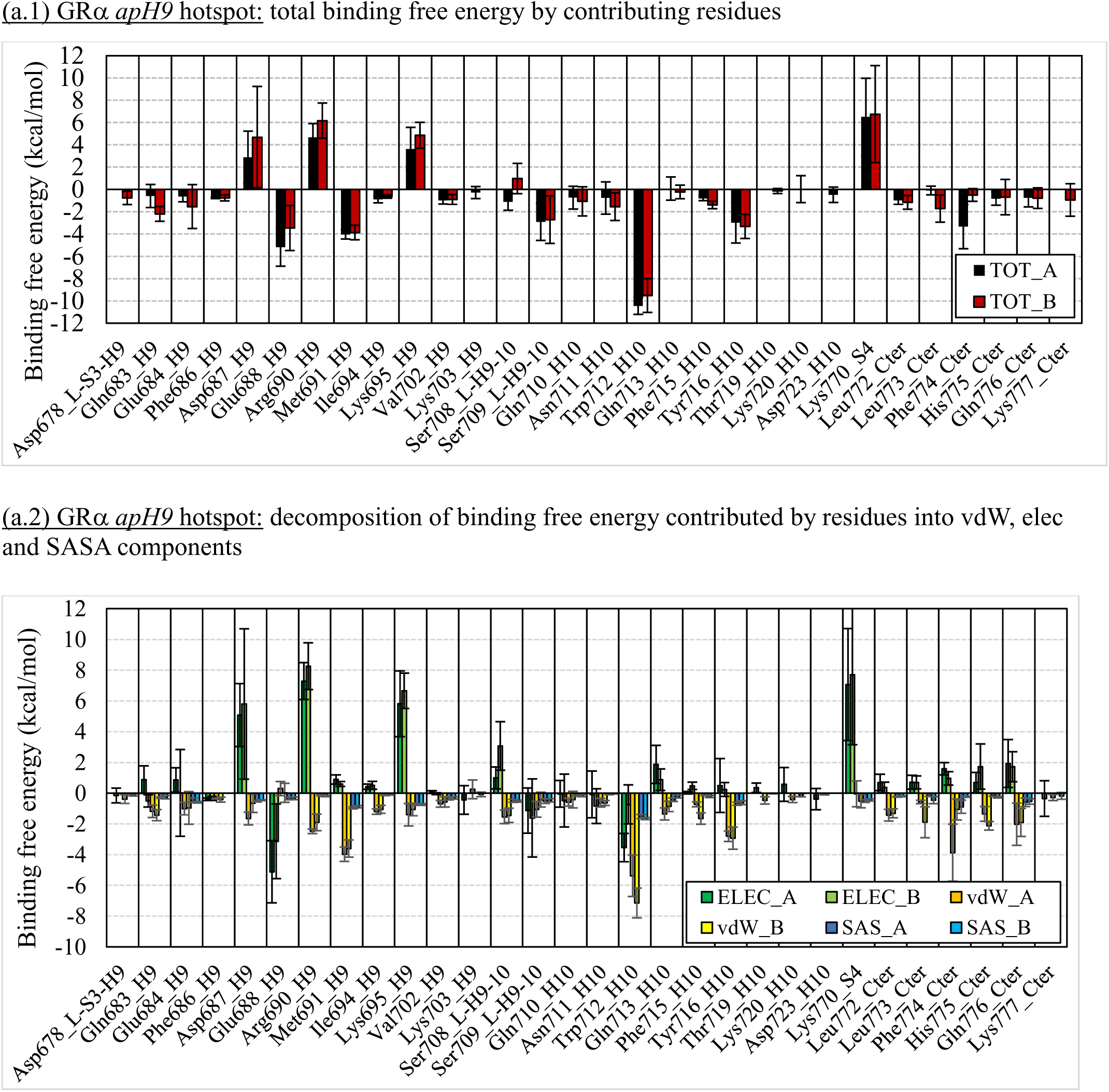
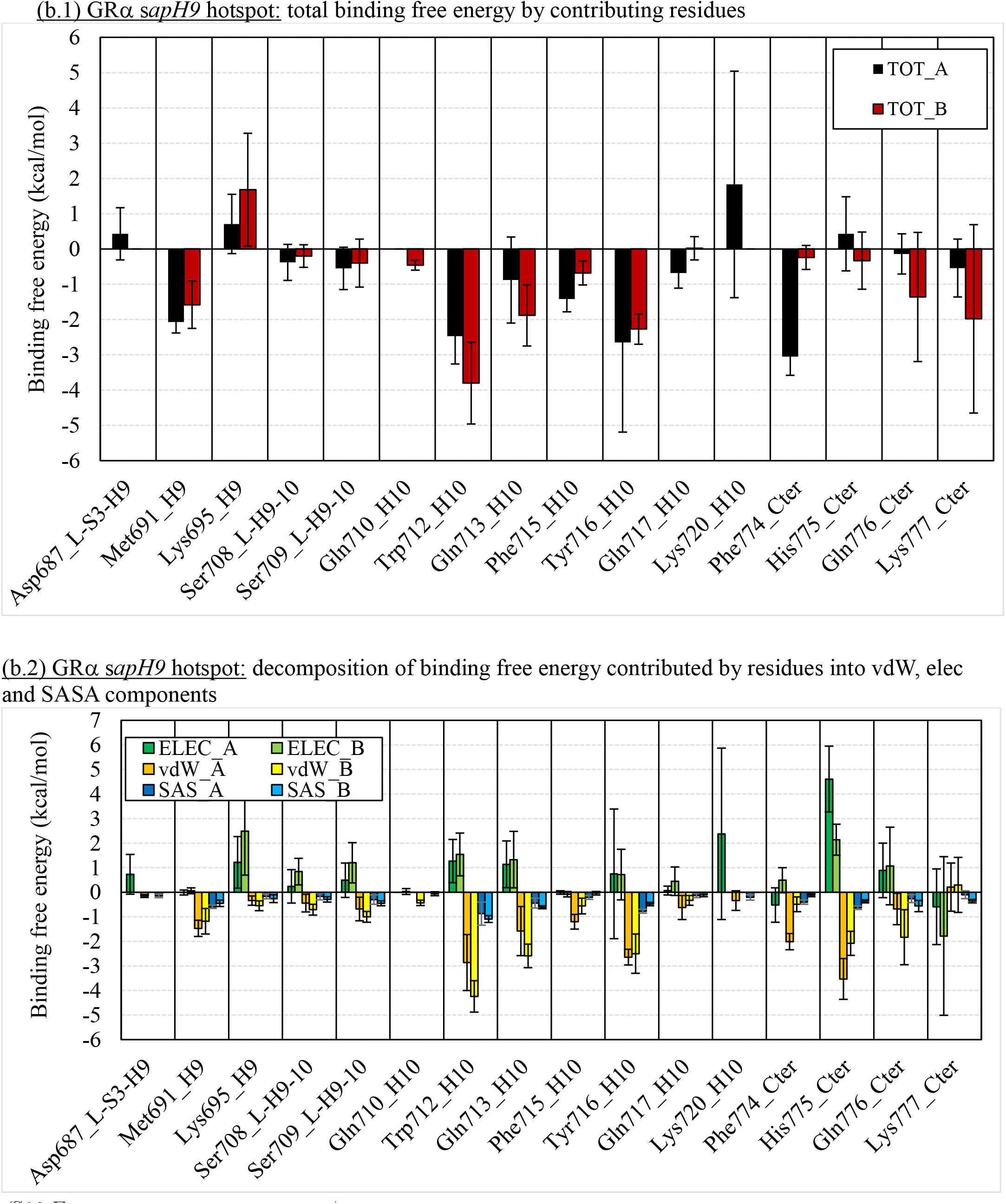
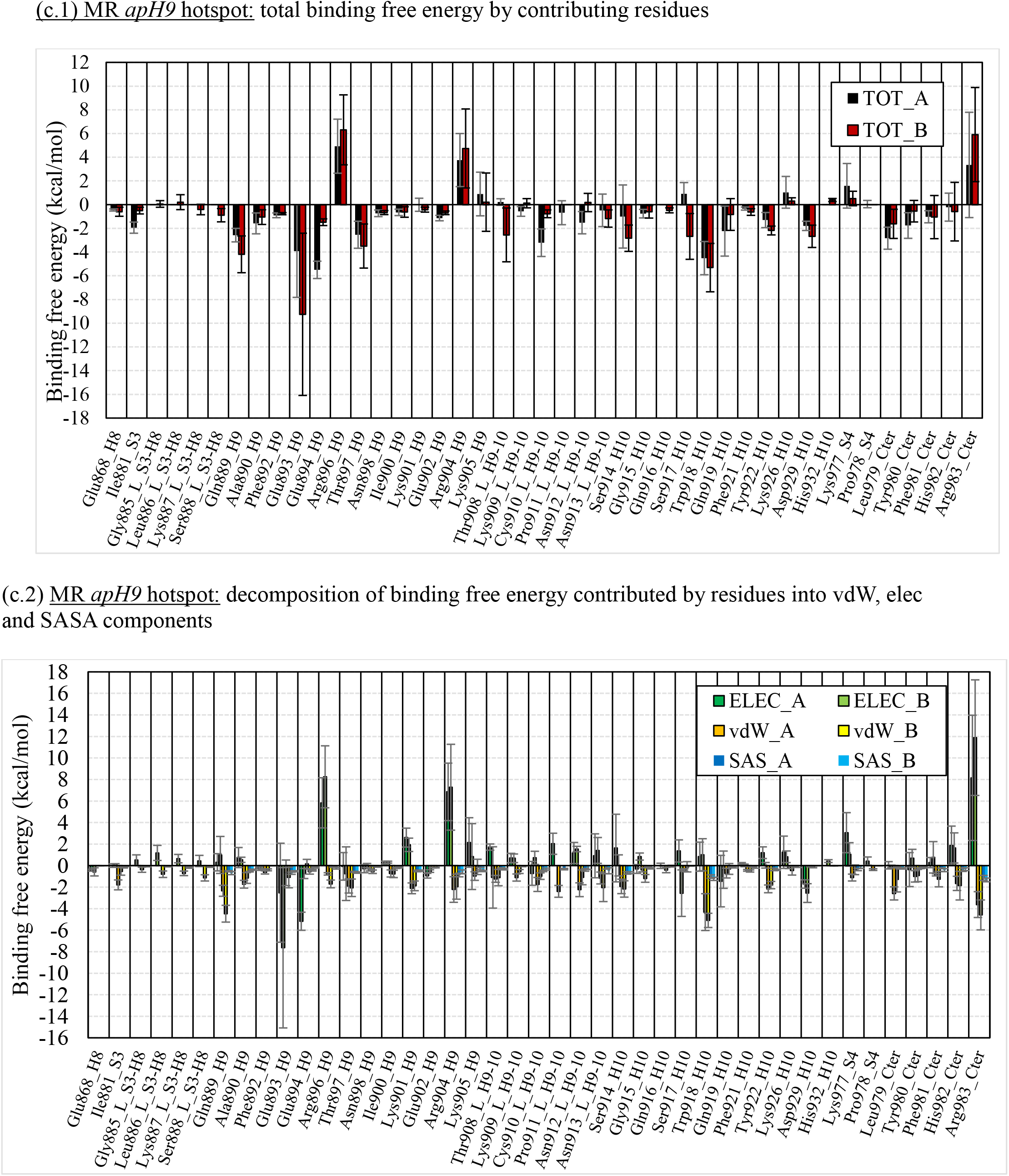
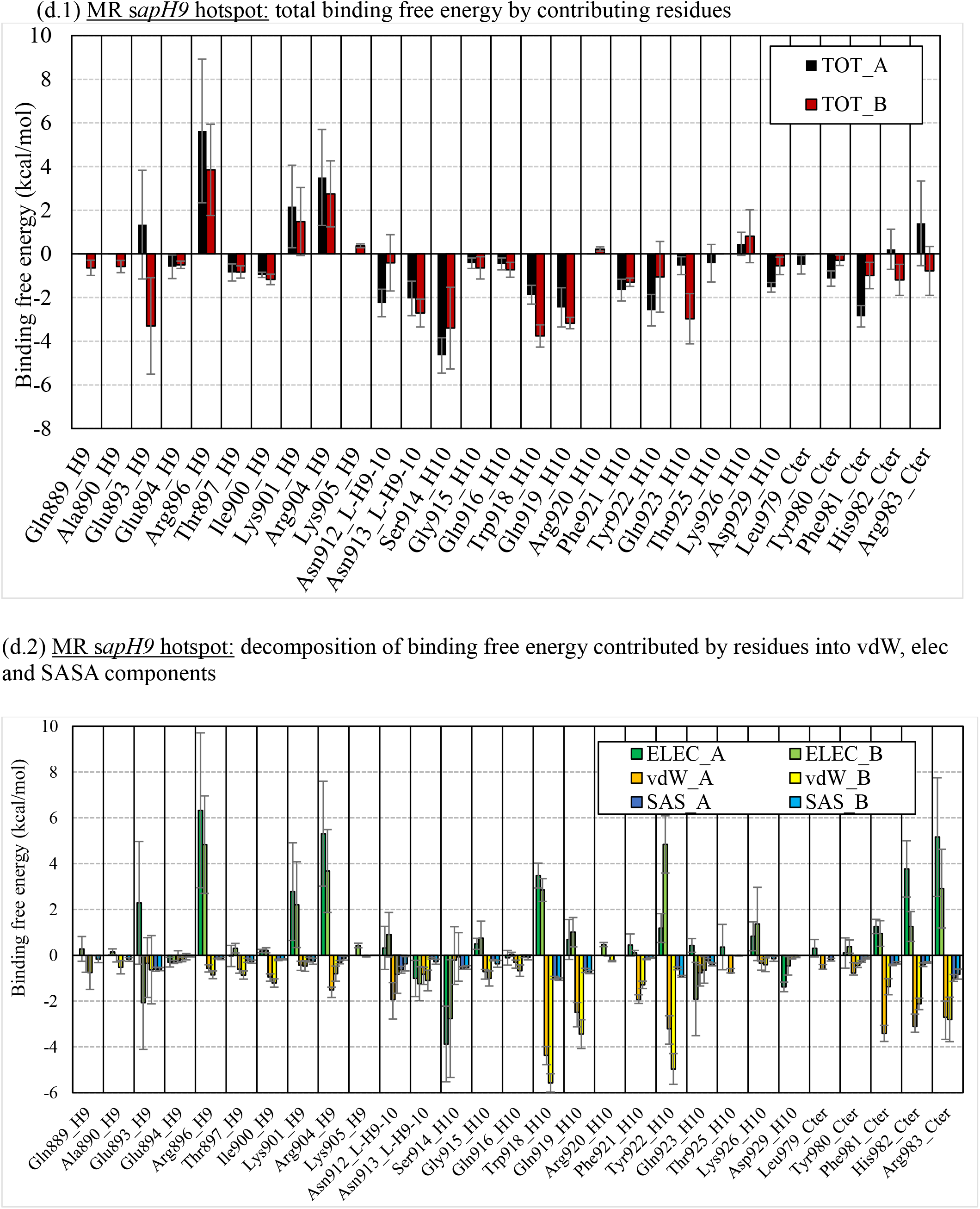
Residues that contribute binding free energy (hotspot) in GRα and MR *sapH9* or *apH9* complexes. PDB-ID:2AAX and PDB-ID:3VHU starting structures were selected for MR *sapH9* and MR *apH9*, respectively while only one starting structure *i.e*. PDB-ID:4P6W was available for GRα. Binding free energies were determined on MD trajectory average structures using the MM/PBSA method.

**S13 Figure:**
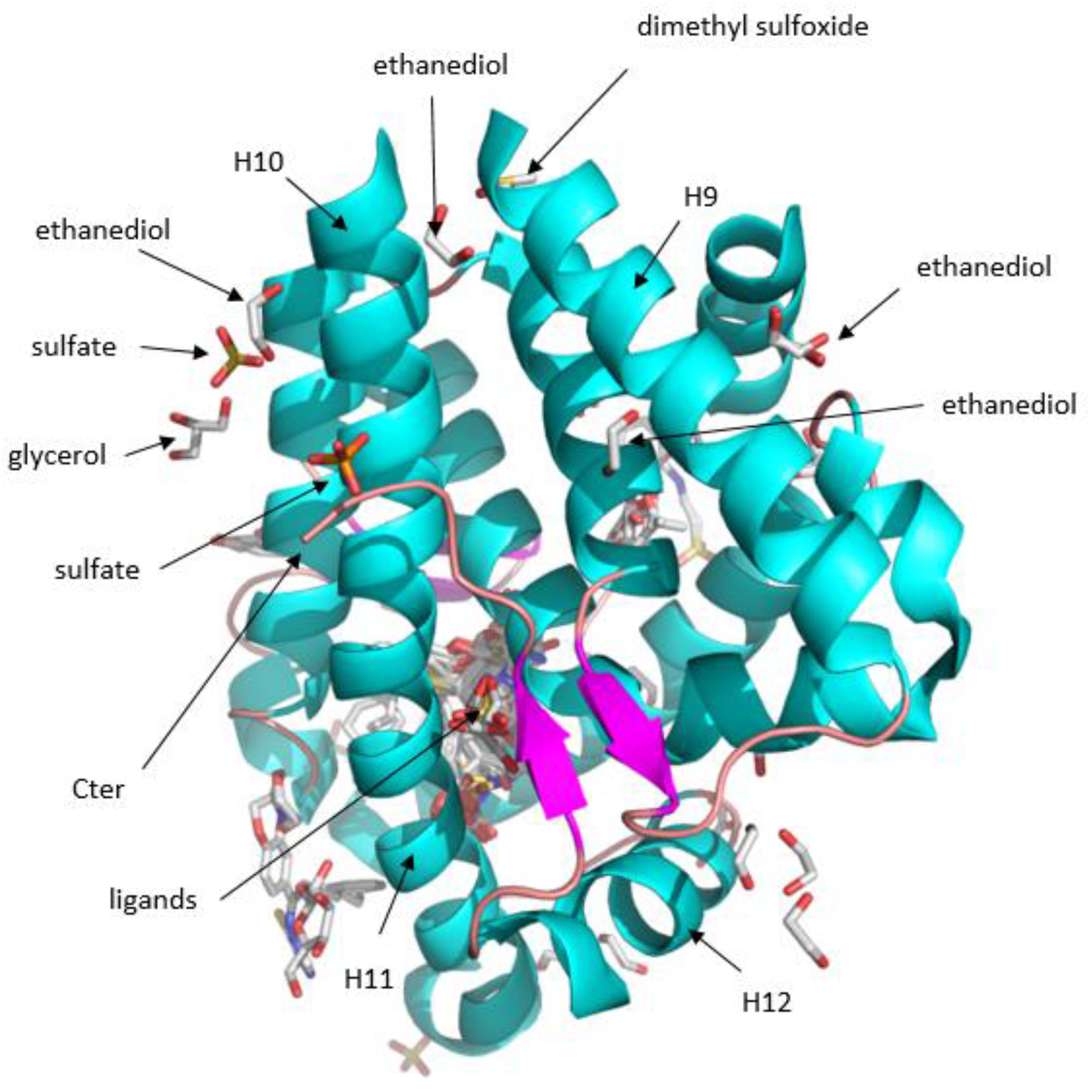
Superimposition of MR LBDs showing both organic and inorganic additives used for protein crystallization.

## Notes

### Competing Interest Statement

The authors have declared no competing interest.

